# Distinct activities of bacterial condensins for chromosome management in *Pseudomonas aeruginosa*

**DOI:** 10.1101/2020.05.18.101659

**Authors:** Virginia S. Lioy, Ivan Junier, Valentine Lagage, Isabelle Vallet, Frédéric Boccard

**Affiliations:** Université Paris-Saclay, CEA, CNRS, Institute for Integrative Biology of the Cell (I2BC), 91198, Gif-sur-Yvette, France; CNRS, Université Grenoble Alpes, TIMC-IMAG, 38000 Grenoble, France; Department of Biochemistry, University of Oxford, Oxford, UK

**Keywords:** Bacterial condensin, Smc, MukBEF, MksBEF, chromosome segregation, chromosome conformation

## Abstract

Bacteria encompass three types of structurally related SMC complexes referred to as condensins. Smc-ScpAB is present in most bacteria while MukBEF is found in enterobacteria and MksBEF is scattered over the phylogenic tree. The contributions of these condensins to chromosome management were characterized in *Pseudomonas aeruginosa* that carries both Smc-ScpAB and MksBEF. In this bacterium, SMC-ScpAB controls chromosome disposition by juxtaposing chromosome arms. In contrast, MksBEF is critical for chromosome segregation in the absence of the main segregation system and affects the higher-order architecture of the chromosome by promoting DNA contacts in the megabase range. Strikingly, our results reveal a prevalence of Smc-ScpAB over MksBEF involving a coordination of their activities with the DNA replication process. They also show that *E. coli* MukBEF can substitute for MksBEF in *P. aeruginosa* while prevailing over Smc-ScpAB. Altogether, our results reveal a hierarchy between activities of bacterial condensins on the same chromosome.

## INTRODUCTION

The genome of every organism must be highly compacted to fit within a cell that is 1000-fold smaller than the size of the DNA molecule itself. Structural features of chromosomes differ largely between eukaryotes and prokaryotes, even though the functional constraints for gene expression and genome stability are the same. Remarkably, Structural Maintenance of Chromosomes (SMC) complexes are molecular machines thought to be capable of remodeling chromosome super-structure in all cells, from bacteria to mammals.

In eukaryotes, three distinct SMC complexes are found, called cohesin, condensin and Smc5/6. They contribute to different aspects of chromosome organization during different phases of the cell cycle. Cohesins, in addition to their role in *trans*-tethering sister chromatids in G2 phase, are involved in chromosome organization during interphase ; they are responsible for the formation of topologically associated domains (TAD), which are lost rapidly upon entry into prophase in a condensin-dependent manner (Gibcus et al., 2018). Eukaryotic condensins are responsible for mitotic chromosome formation by extensive compaction, whereas Smc5/6 is involved in the DNA damage response (Uhlmann, 2016).

In bacteria, chromosome organization relies on several general processes including macromolecular crowding, DNA supercoiling and DNA folding by binding proteins. This combination of processes modulates the probabilities of DNA contacts at different scales and gives rise to a multilayer structuring of the chromosome (Kleckner et al., 2014; Lioy et al., 2018; Wang et al., 2013). As in eukaryotes, bacterial SMC complexes play critical roles in chromosome organization and segregation. Three different SMC complexes (Smc-ScpAB, MukBEF, MksBEF) sharing a similar overall structure have been identified and called “bacterial condensins”. Smc-ScpAB is the most conserved complex (the Smc protein is homologous to eukaryotic SMCs), found in the majority of bacterial species. In contrast, MukBEF and MksBEF structural similarity with Smc-ScpAB may in part result from convergent evolution (Cobbe and Heck, 2004).

The Smc-ScpAB complex often works together with the ParABS system, which is conserved in many bacterial species and is critical for chromosome segregation (Badrinarayanan et al., 2015). The ParB DNA-binding protein recognizes its target *parS*, present in one or several copies in the Ori region of bacterial chromosomes (Livny et al., 2007). ParB is a CTP hydrolase whose activity is critical for its interaction with *parS* (Jalal et al., 2020; Osorio-Valeriano et al., 2019; Soh et al., 2019). The resulting ParB/*parS* nucleoprotein complex encompasses several kbs of DNA and interacts with the ParA ATPase that drives its segregation (Kawalek et al., 2020). The ParB-*parS* complex also recruits Smc-ScpAB onto the chromosome (Gruber and Errington, 2009; Sullivan et al., 2009). This transient association depends on ATP binding by Smc (Wilhelm et al., 2015). Upon ATP hydrolysis, Smc-ScpAB is released from *parS* sites and relocates to the flanking DNA (Minnen et al., 2016), and its subsequent translocation promotes the juxtaposition of chromosome arms (Le et al., 2013; Marbouty et al., 2015; Wang et al., 2015). The translocation of Smc-ScpAB proceeds rapidly (>50 kb/min) and over long distances (∼ 2 Mb) (Tran et al., 2017; Wang et al., 2017), in an ATPase-dependent manner (Wang et al., 2018). Mutations in Smc-ScpAB lead to defects in chromosome management that vary in different bacteria, often including an increase in anucleate cells (Nolivos and Sherratt, 2014).

In some γ-proteobacteria (including *Enterobacteriales, Vibrionales* and *Pasteurellales*), the Smc-ScpAB complex is replaced by the MukBEF complex. It was identified in *Escherichia coli* thirty years ago, in a screen for mutants producing anucleate cells (Niki et al., 1991). In *E. coli, mukB* mutants can be rescued by mutations that increase DNA gyrase activity and negative supercoiling (Sawitzke and Austin, 2000). How the MukBEF complex is loaded onto the chromosome and how it promotes chromosome segregation remain to be characterized. Recently, using Chromosome Conformation Capture, we showed that MukBEF does not align *E. coli* chromosome arms but instead gives rise to long-range *cis*-contacts within replication arms. These contacts are absent in the 800-kb long Ter region, where MatP (the factor responsible for Ter structuring (Mercier et al., 2008)) prevents MukBEF activity (Lioy et al., 2018; Mäkelä and Sherratt, 2020; Nolivos et al., 2016). Interestingly, MukBEF and MatP both belong to a group of proteins that coevolved with the Dam methylase (Brézellec et al., 2006).

The MksBEF complex was identified as distantly related to MukBEF and found scattered over the phylogenic tree (Petrushenko et al., 2011), which questioned the long-standing assumption that bacterial genomes contain only one bacterial condensin involved in chromosome organization, either MukBEF or Smc-ScpAB. Interestingly, all *Pseudomonas* species encode both a Smc-ScpAB and at least one MksBEF complex. This suggests that in contrast to other bacteria, members of the *Pseudomonas* genus have retained two different bacterial condensins during extensive periods of evolution. Thus, these species represent a perfect model to address the questions of whether the presence of several condensins reflects a requirement for multiple chromosome management activities, and how these activities might be coordinated.

We previously showed that the *P. aeruginosa* chromosome is globally oriented from the old pole of the cell to the division plane/new pole along the *oriC*-*dif* axis, and that its Ori region is positioned at the 0.2/0.8 relative cell length in a ParABS-dependent manner (Vallet-Gely and Boccard, 2013). Four *parS* sites located within 15 kb from *oriC* are recognized by ParB; yet, a single *parS* is sufficient for efficient chromosome segregation, depending on its distance from *oriC* (Lagage et al., 2016). Here, using a combination of Chromosome Conformation Capture (3C) methods and fluorescence microscopy, we analyzed the relative contribution of the two *P. aeruginosa* condensins to chromosome management. Our results indicate that MksBEF can extend the range of *cis* contacts of chromosomal loci and support chromosome segregation in the absence of a functional ParABS system, while Smc-ScpAB mainly controls chromosome disposition inside the cell by juxtaposing chromosome arms from *parS*. Because MksBEF activities are reminiscent of MukBEF activities in *E. coli*, we also introduced *E. coli* MukBEF in *P. aeruginosa* to explore its activity in a heterologous host. We show that MukBEF can substitute for MksBEF, but that it prevents Smc-ScpAB from aligning chromosome arms, whereas MksBEF action on chromosome conformation appears to be restricted by Smc-ScpAB. Altogether, our results reveal a hierarchy between the different activities of bacterial condensins on the same chromosome.

## RESULTS

### MksBEF is critical for chromosome segregation in the absence of the ParABS system

The relative contribution to chromosome segregation of the two *Pseudomonas* condensins was analyzed by measuring the amount of anucleate cells present in liquid cultures under slow growth conditions (Figure 1A). This amount reaches 3% for a Δ*parS* mutant whereas it remains very low for mutants deprived of either one of the bacterial condensins (below 0.15%). In the mutant deprived of both condensins, this amount remains considerably lower than in the Δ*parS* mutant (0.5% versus 3%), indicating that the ParABS system plays a preponderant role in *P. aeruginosa* chromosome segregation. Strikingly, the amount of anucleate cells for the Δ*parS* Δ*mks* mutant reaches 24% (8 times the amount of the Δ*parS* mutant), whereas the one for the Δ*parS* Δ*smc* mutant is only slightly increased compared to the Δ*parS* mutant (1.3 times, to 4%). This demonstrates that the MksBEF contribution to the segregation process becomes critical in the absence of a functional ParABS system, in contrast to the Smc-ScpAB contribution. Similar results were obtained in faster growing conditions (Figure S1A).

**Figure 1:**
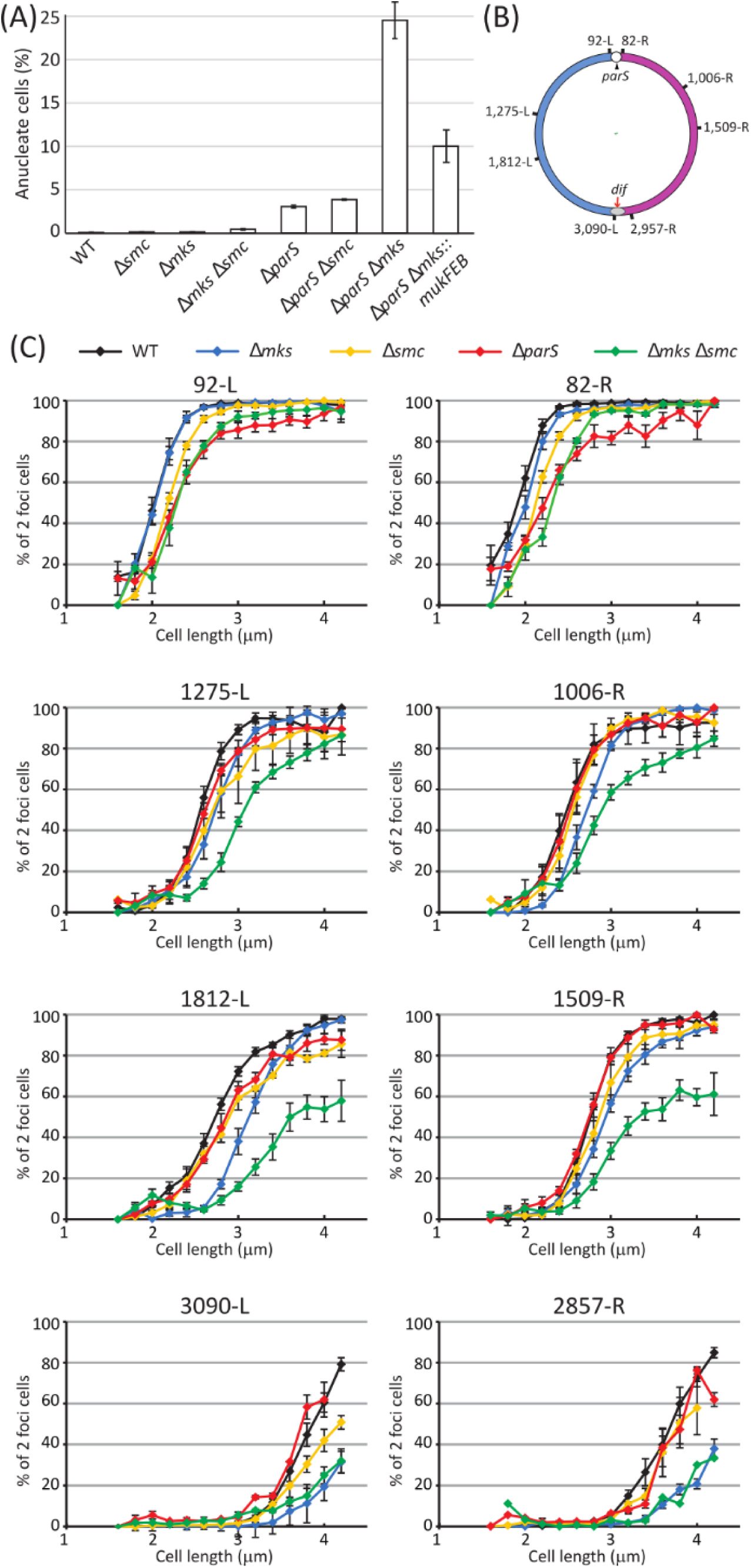
Differential impact of ParABS, MksBEF and Smc-ScpAB on chromosome segregation and separation of chromosomal markers. **(A)** Percentage of anucleate cells (white bars) of the different mutants grown in minimal medium supplemented with citrate at 30°C. Histograms and error bars represent the mean and standard deviation for at least three independent experiments. **(B)** Schematic representation of the position of the chromosomal loci whose duplication and localization inside the cell are studied in **(C).** *oriC* is represented as a white circle, next to the *parS* sites (black arrow). The grey oval represents the terminus of replication and the *dif* site is indicated by a red arrow. *oriC* and *dif* define the left and right chromosome arms (blue and pink, respectively). **(C)** Percentage of two-foci cells according to cell size, in bacterial population grown in minimal medium supplemented with citrate at 30°C (as in every microscopy experiment shown hereafter). Different chromosomal loci (indicated above each graph) are analyzed in different genetic backgrounds: the wild type strain (black), the Δ*mks* mutant (blue), the Δ*smc* mutant (yellow), the Δ*parS* mutant (red) and the Δ*mks* Δ*smc* mutant (green). Values from at least three experiments were averaged, and the error bars represent standard deviations.

### MksBEF and Smc-ScpAB facilitate bulk chromosome separation upon replication, but do not affect the Ori region positioning inside the cell

The dynamics of chromosomal loci separation following replication was also analyzed in the different mutants. Using a snapshot analysis, we represented the amount of two foci cells as a function of cell size to visualize the dynamics of separation of chromosomal loci during the cell cycle (Figure 1B, 1C). The absence of the *parS* sites mostly affects loci located in the Ori region (92-L and 82-R), and not loci located in the Ter region (3090-L and 2857-R). A slight delay is observed for loci of the left chromosome arm (1275-L and 1812-L), but not for loci of the right chromosome arm (1006-R and 1509-R), whose dynamics of separation are identical to those of the wild type strain. In contrast, the absence of the MksBEF condensin has a more pronounced effect for loci located at increasing distance from the *parS* sites, culminating for loci of the Ter region (3090-L and 2857-R). The absence of the Smc-ScpAB condensin induces at most a modest delay in the separation of all loci. However, in the absence of both condensins, a strong delay in separation of loci located in the chromosome arms is observed (1275-L, 1006-R, 1812-L and 1509-R). Considering that the Δ*mks* Δ*smc* mutation generates only a few anucleated cells (Figure 1A), this indicates that the delay in the separation of chromosomal loci does not trigger a strong defect in overall chromosome segregation. Finally, we also recorded the positioning inside the cell of a chromosomal locus close to *oriC* (82-R) in the different mutants (Figure S1B). The two copies of the locus are positioned near the 0.2/0.8 relative cell length in the wild type strain and in the condensin mutants, but not the Δ*parS* mutant. This indicate that the positioning of the Ori region at the 0.2/0.8 relative cell length is only dependent on the ParABS system and not on the condensins. Altogether, this microscopy analysis reveals distinct roles of the ParABS system and the two *P. aeruginosa* condensins during chromosome segregation. It also indicates that both condensins play a role in the separation of loci located farther from the *parS* sites (i.e. in the distal part of the chromosome).

### *P. aeruginosa* chromosome conformation is consistent with its longitudinal disposition inside the cell

Using chromosome conformation capture coupled with deep sequencing (3C-seq (Lioy et al., 2018; Marbouty et al., 2015)), the conformation of the 6.28 Mb *P. aeruginosa* chromosome was analyzed to determine the contribution of ParABS, MksBEF and Smc-ScpAB to this conformation. Contact maps from at least two independent experiments were established at a 10kb-bin resolution, and are represented with *oriC* positioned in the middle of the axes, with the left and right chromosome arms on either side. The contact map of the wild type strain reveals the presence of two diagonals (Figure 2A and Figure S2A). The predominant one (from top left corner to bottom right corner; called hereafter “the primary diagonal”) represents *cis* contacts between neighboring loci, whereas the orthogonal one (from bottom left corner to top right corner hereafter called “the secondary diagonal”) represents contacts between loci located on the two chromosome arms (it comprises less than 10% of the total amount of contacts). This secondary diagonal is consistent with the longitudinal disposition of the *P. aeruginosa* chromosome observed by microscopy (Vallet-Gely and Boccard, 2013), and reveals a close proximity of the chromosome arms. Chromosome domains of various sizes (reminiscent of the CIDs described in other bacteria (Le et al., 2013; Lioy et al., 2018; Marbouty et al., 2015; Wang et al., 2015) are observed along the primary diagonal. They are mostly conserved in the contact map of the different mutants investigated here, and their systematic detailed analysis is the subject of another study (Varoquaux et al. 2020 in revision, and our unpublished results).

**Figure 2:**
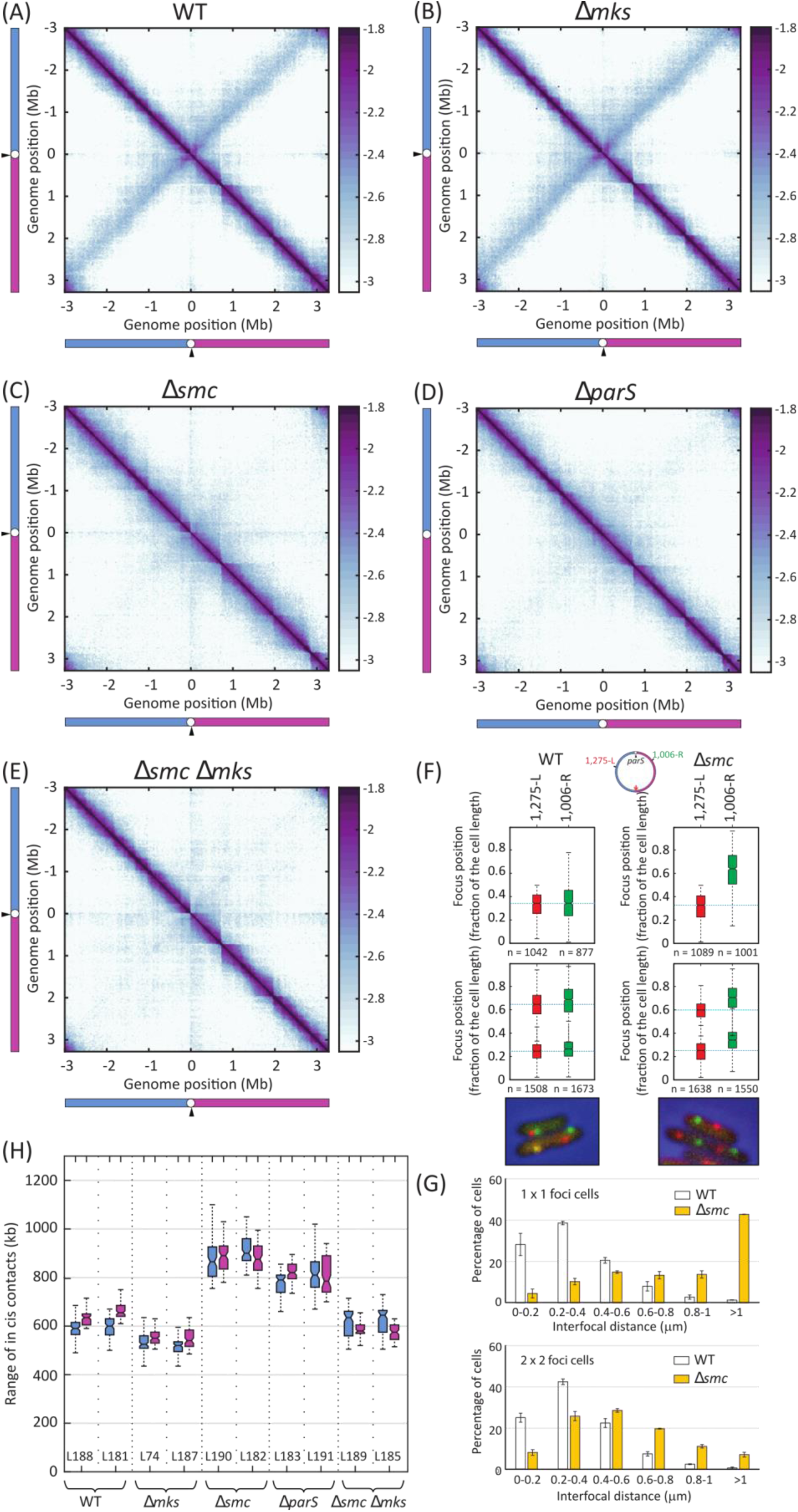
Differential impact of ParABS, MksBEF and Smc-ScpAB on chromosome conformation and disposition inside the cell. **(A-E)** Normalized contact maps obtained for different strains grown in minimal medium supplemented with citrate at 30°C. Abscissa and ordinate axis represent genomic coordinates, *oriC* being located in the middle (white circle), next to the *parS* sites (black arrow), with the left and right chromosome arms indicated in blue and pink, respectively. The same conventions will be used hereafter to schematize chromosome maps of the different strains. The color scale reflects the frequency of contacts between two regions of the genome, from white (rare contacts) to dark purple (frequent contacts). Maps obtained for the wild type strain **(A)**, the Δ*mks* mutant **(B)**, the Δ*smc* mutant **(C)**, the Δ*parS* mutant **(D)** and the Δ*smc* Δ*mks* mutant **(E). (F)** Relative position inside the cells of two chromosomal loci located at similar distance from *oriC* (highlighted in green and red), in the wild type strain (left panels) and in the Δ*smc* mutant (right panels). Boxplot representations are used, indicating the median (horizontal bar), the 25th and the 75^th^ percentile (open box) and the rest of the population except for the outliers (whiskers). Outliers are defined as 1.5×*IQR* or more above the 75^th^ percentile or 1.5×*IQR* or more below the first 25th percentile quartile. Cells are arbitrarily oriented, the 0 pole being the one closest to the 1,812-L locus. Representative images are shown below the graphs. Experiments have been performed at least twice independently; one representative example is shown here. **(G)** Interfocal distances (in micrometers) between the two chromosomal loci, in cells containing one copy of each locus (top panel) or two copies of each locus (bottom panel). The percentage of cells in which the distance is of a certain value is plotted, for the wild type strain (white) and the Δ*smc* mutant (yellow). Histograms and error bars represent the mean and standard deviation for two independent experiments. **(H)** Quantification of the range of *cis* interactions of chromosomal loci located on the left chromosome arm (in blue) or on the right chromosome arm (in pink) of strains presenting different genetic backgrounds (see supplementary Figure 3 for details). Results are shown for each replicate (contact map numbers are indicated below). Boxplot representations are used, as described above.

### The absence of Smc-ScpAB leads to loss of juxtaposition of *P. aeruginosa* chromosome arms and global repositioning of the chromosome

The contact map obtained from the Δ*mks* mutant presents both diagonals (Figure 2B and S2B). By contrast, in the contact maps obtained from a Δ*smc* or a Δ*parS* mutant, only the primary diagonal is detected (Figure 2C, 2D, S2C and S2D), indicating that chromosome arm proximity depends upon the Smc-ScpAB complex and the *parS* sites, as observed in *C. crescentus* and *B. subtilis* (Le et al., 2013; Marbouty et al., 2015; Wang et al., 2015). In the presence of a single *parS* site, the contact map also presents a secondary diagonal (Figure S2F), indicating that one *parS* site is sufficient for chromosome arm alignment by Smc-ScpAB. In the absence of ParA, the secondary diagonal is also observed in the contact map (Figure S2G), indicating that ParA is not involved in chromosome arm alignment by Smc-ScpAB.

Considering that the absence of Smc-ScpAB provokes a lack of close proximity between the chromosome arms (as indicated by the absence of the secondary diagonal), we explored its consequences for chromosome arm disposition inside the cell by fluorescent microscopy. We compared in the wild type strain and the Δ*smc* mutant the relative position of two different couples of fluorescent tags located at equivalent distances from the *parS* sites, on different chromosome arms (Figure 2F and Figure S1C). As expected, the two tags are located close together in the wild type strain: when one tag is located in one cell half, the second tag is mostly localized in the same cell half. Strikingly, upon Smc inactivation, the second tag is mostly found in the opposite cell half. This is also true in cells carrying duplicated copies of the two loci (i.e. more advanced in the cell cycle): they are mostly localized close together in the wild type strain whereas the Δ*smc* mutant presents an alternation of the different tags. Consistently, interfocal distances between the two tags are dramatically increased in the Δ*smc* mutant compared to the wild type strain (Figure 2G). Altogether, these results indicate that the longitudinal disposition of the chromosome is lost in the absence of Smc-ScpAB.

### MksBEF activity extends the range of *cis* contacts along chromosome arms

When comparing the contact maps of the different mutants, the primary diagonal appears wider in the Δ*smc* and Δ*parS* mutants (Figure 2C, 2D, S2C and S2D) than in the wild type strain, the Δ*mks* mutant or the Δ*smc* Δ*mks* mutant (Figure 2A, 2B, 2E, S2A, S2B and S2E). The width of the primary diagonal corresponds to the range of *cis* contacts of chromosomal loci (i.e. the DNA length at which a chromosomal locus can interact with its neighboring loci). To assess this range, we used the quantification method developed by Wang and colleagues to measure the kinetics of arm alignment by Smc-ScpAB in *B. subtilis* (Wang et al., 2017), and apply it to the *P. aeruginosa* contact maps (for details see Materials and Methods, Figure S3 and S4). Using this method, we determined that the median range of *cis* contacts of a chromosomal locus located on the chromosome arms of the wild type strain is approximatively 600 kb in both directions, whereas it exceeds 850 kb in the Δ*smc* and Δ*parS* mutants (Figure 2H). The inactivation of MksBEF in a Δ*smc* mutant leads to a strong reduction of the range of *cis* contacts, and this range is similar to that of the wild type strain (around 600 kb). Remarkably, no significant difference is observed in the Δ*mks* mutant compared to the wild type strain. Altogether, these results reveal that MksBEF extends the range of *cis* contacts along the chromosome arms when they are not aligned by Smc-ScpAB (either when Smc-ScpAB is absent (Δ*smc* mutant), or when there is no *parS* site to recruit the Smc-ScpAB complex onto the chromosome (Δ*parS* mutant)). This suggests that alignment by Smc-ScpAB limits MksBEF action on the chromosome arms, which might reflect interference between these two different bacterial condensins acting on the same DNA molecule.

### Uncoupling replication initiation and Smc-ScpAB loading triggers MksBEF-dependent extension of the range of *cis* contacts

We then questioned the impact on *P. aeruginosa* chromosome conformation of modifying the chromosome arm alignment, by displacing the Smc-ScpAB entry point onto the chromosome. To do that, we used two different strains containing a single *parS* site located either 550 kb away from *oriC* on the right chromosome arm (the *parS*^+550^ strain), or 330 kb away from *oriC* on the left chromosome arm (the *parS*^-330^ strain). As previously mentioned, one single *parS* site is sufficient for chromosome arm alignment by Smc-ScpAB. The contact maps of these strains (Figure 3A, 3D, S2J and S2M) indicate that the secondary diagonal is displaced according to the *parS* location on the genome. It appears more variable in intensity than in the wild type strain (the frequency of contacts between the two chromosome arms is higher near the *parS* site than in the distal part), however the entirety of the chromosome arms are juxtaposed.

**Figure 3:**
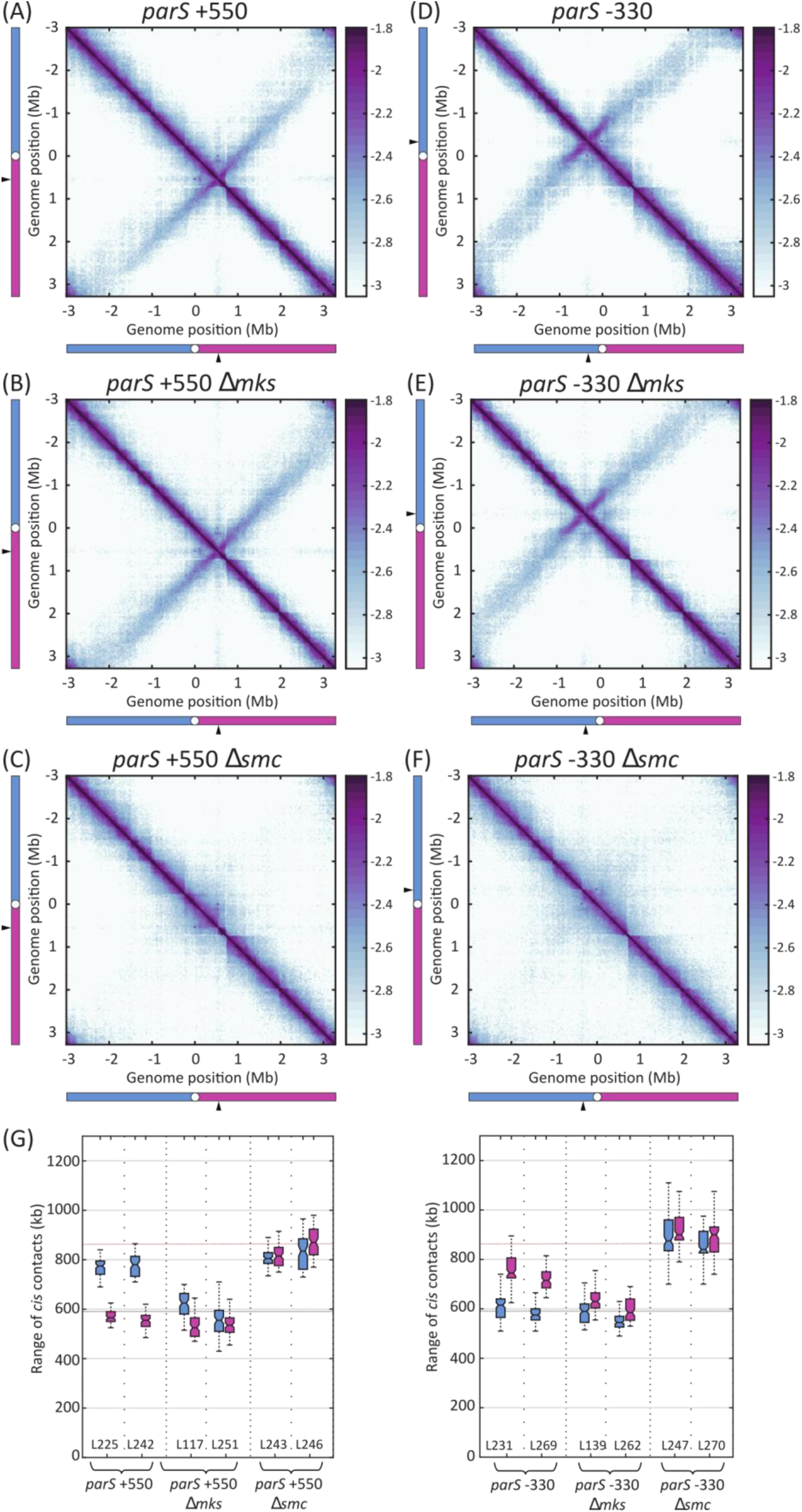
Impact of displacing the *parS* site away from *oriC* on *P. aeruginosa* chromosomal conformation. Normalized contact maps obtained for the *parS* ^+550^ **(A)**, *parS* ^+550^ Δ*mks* **(B)**, *parS*^+550^ Δ*smc* **(C)**, *parS*^-330^ **(D)**, *parS*^-330^ Δ*smc* **(E)**, *parS*^-330^ Δ*mks* **(F)** strains. Abscissa and ordinate axis represent genomic coordinates. The color scale reflects the frequency of contacts between two regions of the genome, from white (rare contacts) to dark purple (frequent contacts). **(G)** Quantification of the range of *cis* interactions of chromosomal loci located on the left chromosome arm (in blue) or on the right chromosome arm (in pink). The horizontal black dotted line indicates the median of *cis* contacts for chromosomal loci belonging to the left arm of the wild type strain (Figure 2A), whereas the horizontal red dotted line indicates the median of *cis* contacts for chromosomal loci belonging to the left arm of the Δ*smc* mutant (Figure 2C). Results are shown for each replicate (contact map numbers are indicated below).

Another striking feature of these contact maps is the difference observed in the range of *cis* contacts for loci belonging to the two chromosome arms (Figure 3G). This range is extended for the chromosome arm that does not encompass the *parS* site (median around 770 kb for the left arm of the *parS*^+550^ strain (in which *parS* is displaced onto the right arm) and around 730 kb for the right arm for the *parS*^-330^ strain (in which *parS* is displaced onto the left arm). In contrast, the median range of *cis* contacts for the other arm is similar to the wild type strain, around 600 kb. Remarkably, in the absence of MksBEF (in the *parS*^+550^ Δ*mks* and the *parS*^-330^ Δ*mks* strains, Figure 3B, 3E, 3G, S2K and S2N), no extension of the range of *cis* contacts is observed in either arm. In contrast, in the absence of Smc-ScpAB (in the *parS*^+550^ Δ*smc* and the *parS*^-330^ Δ*smc* strains, Figure 3C, 3F, 3G, S2L and S2O), an extension of the median range of *cis* contacts is observed for chromosomal loci belonging to both arms (up to 900 kb).

Altogether, these results demonstrate that when the *parS* site is displaced away from *oriC*, MksBEF can extend the range of *cis* contacts for chromosomal loci belonging to the chromosome arm that does not encompass the displaced *parS* site. Considering the relative position of *oriC* and *parS*, replication initiation at *oriC*, i.e. prior to the replication of the *parS* site, creates a delay between the replication of chromosomal loci belonging to this arm and their alignment by Smc-ScpAB. Our data strongly suggest that this delay is necessary for the MksBEF-dependent extension of the range of *cis* contacts. In other words, MksBEF extends the range of *cis* contacts of chromosomal loci when it can act prior to Smc-ScpAB-dependent juxtaposition of chromosome arms.

### MksBEF-dependent extension of the range of *cis* contacts is dependent upon replication initiation

To test this hypothesis further, we used a second strategy, which required the engineering of a strain carrying two origins of replication, the native *oriC* located close to the *parS* sites and an ectopic *oriC* located 923 kb away in the right chromosome arm (called the *oriC ins1* strain). We checked that the ectopic origin was indeed functional by Marker Frequency Analysis (Figure S5), meaning that newly replicated regions are generated from both *oriC*. Considering that chromosome arm alignment proceeds from the *parS* sites, located next to the native *oriC*, there should be a delay between the replication of the DNA regions surrounding the ectopic *oriC* and their alignment by Smc-ScpAB. Strikingly, the contact map of this strain carrying two *oriC* shows that the median range of *cis* contacts of chromosomal loci belonging to the right chromosome arm (containing the ectopic *oriC*) is extended to almost 800 kb, which is not the case for the left chromosome arm (it remains close to 600 kb, Figure 4A, 4D and S2P). This is due to MksBEF, as this extension of the range of *cis* contacts for the right chromosome arm is not observed in the absence of MksBEF (Figure 4B, 4D and S2Q). In contrast, in the absence of Smc-ScpAB, both chromosome arms present an extended median range of *cis* contacts to more than 850 kb (Figure 4C, 4D and Figure S2R), attesting that in the absence of Smc-ScpAB, MksBEF can act on both chromosome arms. Altogether, these results confirm that introducing an artificial delay between replication and alignment of chromosomal regions allow MksBEF to extend the range of *cis* contacts in these regions. They also strongly suggest that MksBEF acts on newly replicated DNA regions.

**Figure 4:**
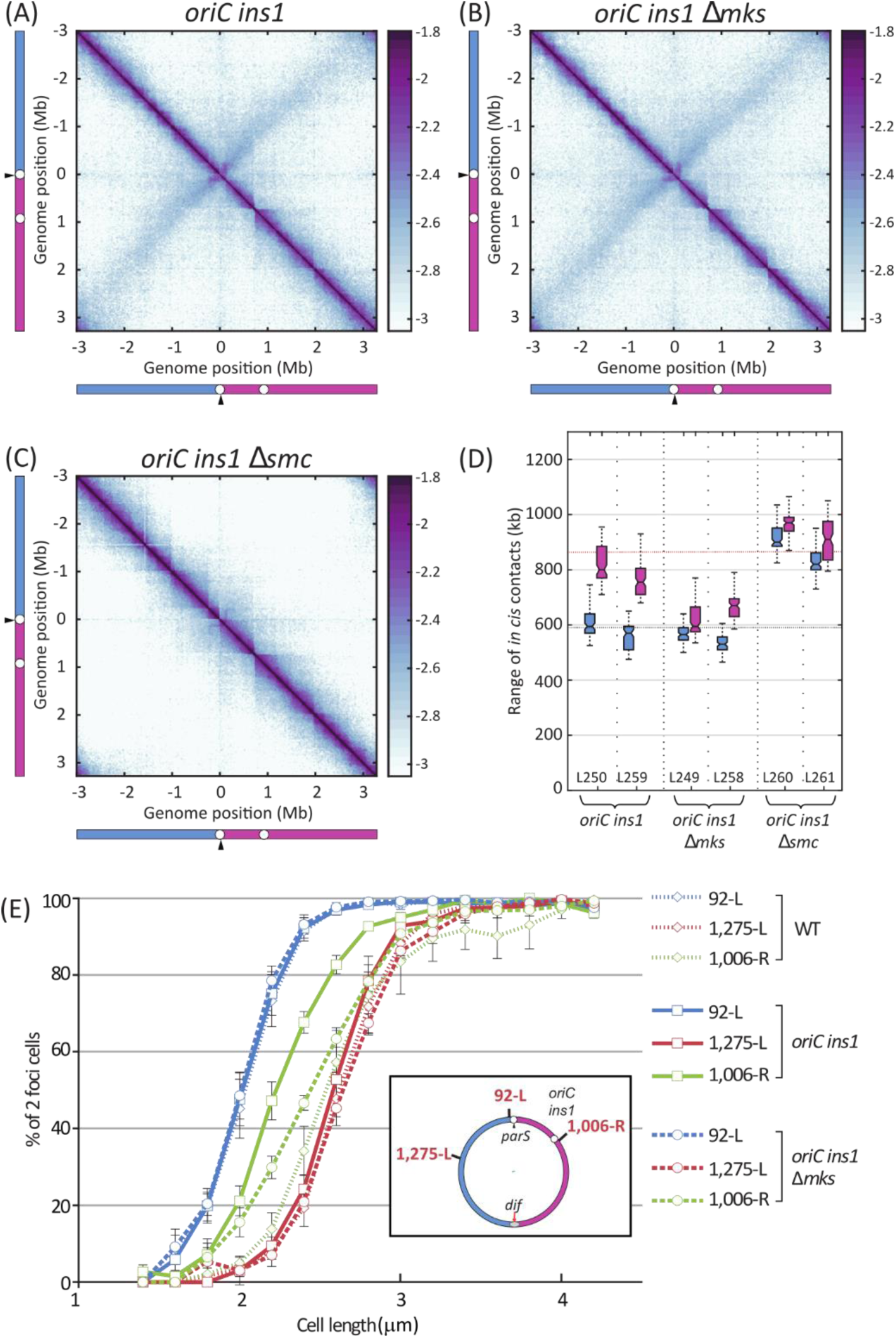
Impact of an additional ectopic *oriC* on *P. aeruginosa* chromosomal conformation and segregation. Normalized contact maps obtained for the *oriC ins1* **(A)**, *oriC ins1* Δ*mks* **(B)** and *oriC ins1* Δ*smc* **(C)** strains. Abscissa and ordinate axis represent genomic coordinates. The color scale reflects the frequency of contacts between two regions of the genome, from white (rare contacts) to dark purple (frequent contacts). **(D)** Quantification of the range of *cis* interactions of chromosomal loci located on the left chromosome arm (in blue) or on the right chromosome arm (in pink). The horizontal black and red dotted lines are as in Figure 3. Results are shown for each replicate (contact maps numbers are indicated below). **(E)** Percentage of two-foci cells according to cell size, in bacterial population grown in minimal medium supplemented with citrate at 30°C, for different fluorescent tags (92-L in blue), 1275-L in red and 1006-R in green) in different genetic backgrounds (in the wild type strain in short-dashed lines, in *oriC ins1* in full lines and in *oriC ins1* Δ*mks* in long-dashed lines). Cells containing the 92-L and one of the other tags were analyzed. The amount of two-foci cells is represented according to cell length.

### Segregation of newly replicated region by MksBEF

To determine whether MksBEF action on newly replicated regions might affect their dynamics of separation during the cell cycle, we used fluorescent microscopy to measure the proportion of two foci cells according to cell size for three chromosomal loci, one located near the native *oriC* (92-L), one located near the insertion site of the ectopic *oriC* (1006-R), and one located on the other chromosome arm (1275-L). We compared the results obtained in the engineered strains carrying two *oriC* (in the presence and in the absence of MksBEF) to those obtained in the wild type strain (Figure 4E). In the wild type strain, the chromosomal locus located close to *oriC* is separated earlier than the two other loci, located farther away on the chromosome arms. Strikingly, in the strain carrying two *oriC*, the separation of the locus located next to the ectopic *oriC* (1006-R) is advanced compared to the wild type strain. This is neither the case for the locus located on the other chromosome arm (1275-L), nor for the locus close to the native *oriC* (92-L). Finally, comparing the dynamics of separation of the three loci in the strains carrying two *oriC* in the presence and in the absence of MksBEF show that the separation of the control loci (92-L and 1,275-L) remains unchanged, whereas the separation of the locus next to the ectopic *oriC* (1006-R) is delayed in the absence of MksBEF. Altogether, these results indicate that MksBEF is indeed able to act on newly replicated regions, and that this action leads to early separation of chromosomal loci.

### Analogous impact of MksBEF and MukBEF on the *P. aeruginosa* chromosome

The results presented above indicate that (i) MksBEF is critical for chromosome segregation in the absence of ParABS, and that (ii) MksBEF can extend the range of *cis* contacts along chromosome arms. These two features are reminiscent of the role of MukBEF in *E. coli* (that lacks a ParABS system), which is particularly striking considering that MksBEF was initially identified as distantly related to MukBEF. To investigate an eventual functional relationship between these two condensins, we replaced the MksBEF coding sequences by the MukBEF coding sequences in the *P. aeruginosa* genome, and we analyzed the impact of this replacement on chromosome segregation and chromosome conformation.

Anucleate cell measurements (Figure 1A and S1A) shows that the amount of anucleate cells for the Δ*parS* Δ*mks* mutant expressing the *mukFEB* genes is lower than for the Δ*parS* Δ*mks* mutant in slow growing conditions. They also show that MukBEF can complement the absence of MksBEF in faster growing conditions (Figure S1A). Furthermore, as observed with MksBEF, in a strain carrying two *oriC*, MukBEF is able to act on newly replicated regions leading to early separation of chromosomal loci (Figure 5A). Altogether, this indicates that MukBEF is able to promote chromosome segregation in *P. aeruginosa* in the absence of a functional ParABS system.

**Figure 5:**
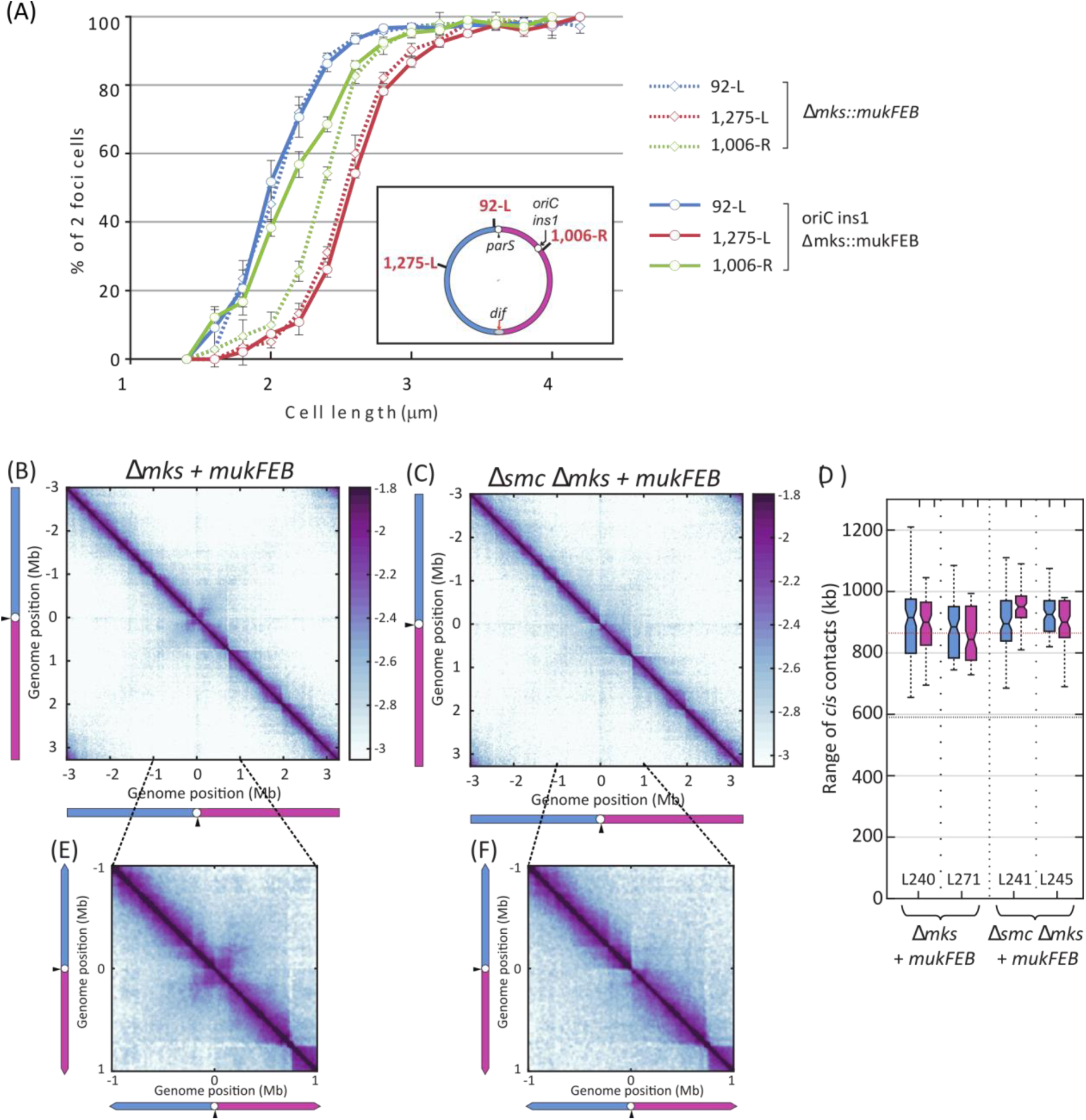
MukBEF impact on *P. aeruginosa* chromosome conformation. **(A)** Percentage of two-foci cells according to cell size, in bacterial populations grown in minimal medium supplemented with citrate at 30°C, for different fluorescent tags (92-L in blue, 1275-L in red and 1006-R in green) in different genetic backgrounds (in the Δ*mks*::*mukBEF* strain in dashed lines, and in the *oriC ins1* Δ*mks*::*mukBEF* strain in full lines). Cells containing the 92-L and one of the other tags were analyzed. The percentage of two-foci cells is represented according to cell length. **(B, C)** Normalized contact maps obtained for the Δ*mks*::*mukFEB* **(B)** and the Δ*smc* Δ*mks*::*mukFEB* strains **(C)**. Abscissa and ordinate axis represent genomic coordinates. The color scale reflects the frequency of contacts between two regions of the genome, from white (rare contacts) to dark purple (frequent contacts). **(D)** Quantification of the range of *cis* interactions of chromosomal loci located on the left chromosome arm (in blue) or on the right chromosome arm (in pink) of strains in which the *mksFEB* operon has been replaced by the *mukFEB* operon. The horizontal black and red dotted line are as in Figure 3. Results are shown for each replicate (contact map numbers are indicated below). **(E)** and **(F)** Zoom of the contact maps on the 2 Mb region surrounding *oriC*, for the Δ*mks::MukFEB* strain and the Δ*smc* Δ*mks::MukFEB* mutant respectively.

Using 3-C seq, we also analyzed chromosome conformation of strains expressing *mukFEB* instead of *mksFEB*. Only the primary diagonal is observed on contact maps established for the Δ*mks* and Δ*smc* Δ*mks* mutants expressing the *mukFEB* genes (Figure 5B, 5C, S2S and S2T). This reveals that chromosome arm alignment by Smc-ScpAB does not occur in the presence of MukBEF. Moreover, the range of *cis* contacts along chromosome arms is systematically extended when MukBEF is expressed (Figure 5D, to more than 900 kb), which demonstrates that MukBEF can enhance the range of *cis* contacts in *P. aeruginosa* whether Smc-ScpAB is present or not.

Altogether, these results indicate that MksBEF and MukBEF are both able to promote chromosome segregation in the absence of a functional ParABS system and to extend the range of *cis* contacts of chromosomal loci in *P. aeruginosa*. However, their functional interplay with Smc-ScpAB differs strikingly, as MukBEF prevents chromosome arm alignment by Smc-ScpAB whereas MksBEF action appears to be restricted by Smc-ScpAB.

## DISCUSSION

### Two bacterial condensins in *P. aeruginosa* present distinct activities for chromosome conformation

Most bacteria present a single condensin complex, either Smc-ScpAB or MukBEF. Thus, *Pseudomonas* species constitute a remarkable genus, as two different complexes, Smc-ScpAB and MksBEF, are both involved in chromosome management. We show here that these two complexes present distinct activities in *P. aeruginosa*, with Smc-ScpAB promoting chromosome arms alignment and controlling chromosome disposition inside the cell, while MksBEF promotes the extension of the range of *cis* contacts in the absence of chromosome arm alignment, and contributes to chromosome segregation in the absence of the ParABS system. Remarkably, among the >100 *Pseudomonas* species whose genome sequence is known, all of them encode both condensins, revealing a fixation of both condensin activities since the divergence of the *Pseudomonas* genus (more than 1 billion years ago, (Battistuzzi et al., 2004)). It is interesting to note that different bacterial species including some *Pseudomonas* strains encode another Mks complex called MksBEFG containing a fourth component (Petrushenko et al., 2011), which seems to be involved in plasmid maintenance rather than in chromosome management (Doron et al., 2018; Panas et al, 2014; Bohm et al., 2020). That MksBEF and MksBEFG complexes share a common ancestor is unlikely, as they do not show sequence similarity. Further investigations will be needed to understand the evolution of this large family of proteins.

We demonstrate that there is a hierarchy of condensin activities, as MksBEF-dependent extension of the range of *cis* contacts was only revealed in the absence of chromosome arm alignment by Smc-ScpAB in the wild type genome configuration (Figure 2). In this configuration, *parS* is located close to *oriC* (less than 15kb) ensuring that chromosome arm alignment by Smc-ScpAB starts quickly following replication initiation. We propose that this synchronicity between replication and alignment by Smc-ScpAB of chromosomal loci prevents MksBEF dependent extension of their range of *cis* contacts. In support of this hypothesis, we show that engineering new genome configurations by displacing the *parS* sites away from *oriC* or adding an ectopic *oriC* far from the *parS* sites allow the detection of both Smc-ScpAB and MksBEF activities on the same chromosome (Figures 3, 4 and Figure S6). As Smc-ScpAB chromosome arm alignment initiates at *parS* upon its replication, these engineered configurations lead to a delay between the replication of certain chromosome regions and their alignment by Smc-ScpAB. This delay allows the detection of the MksBEF-dependent extension of the range of *cis* contacts in chromosomal regions replicated long before Smc-ScpAB can align them. Altogether, these results support a model whereby Smc-ScpAB aligns the replication arms upon replication of the *parS* sites whereas MksBEF gives rise to long-range *cis* contacts in newly replicated regions provided MksBEF can act before Smc-ScpAB. Taken together, this suggests coordination of both condensin activities with the replication process (Figure 6A).

**Figure 6:**
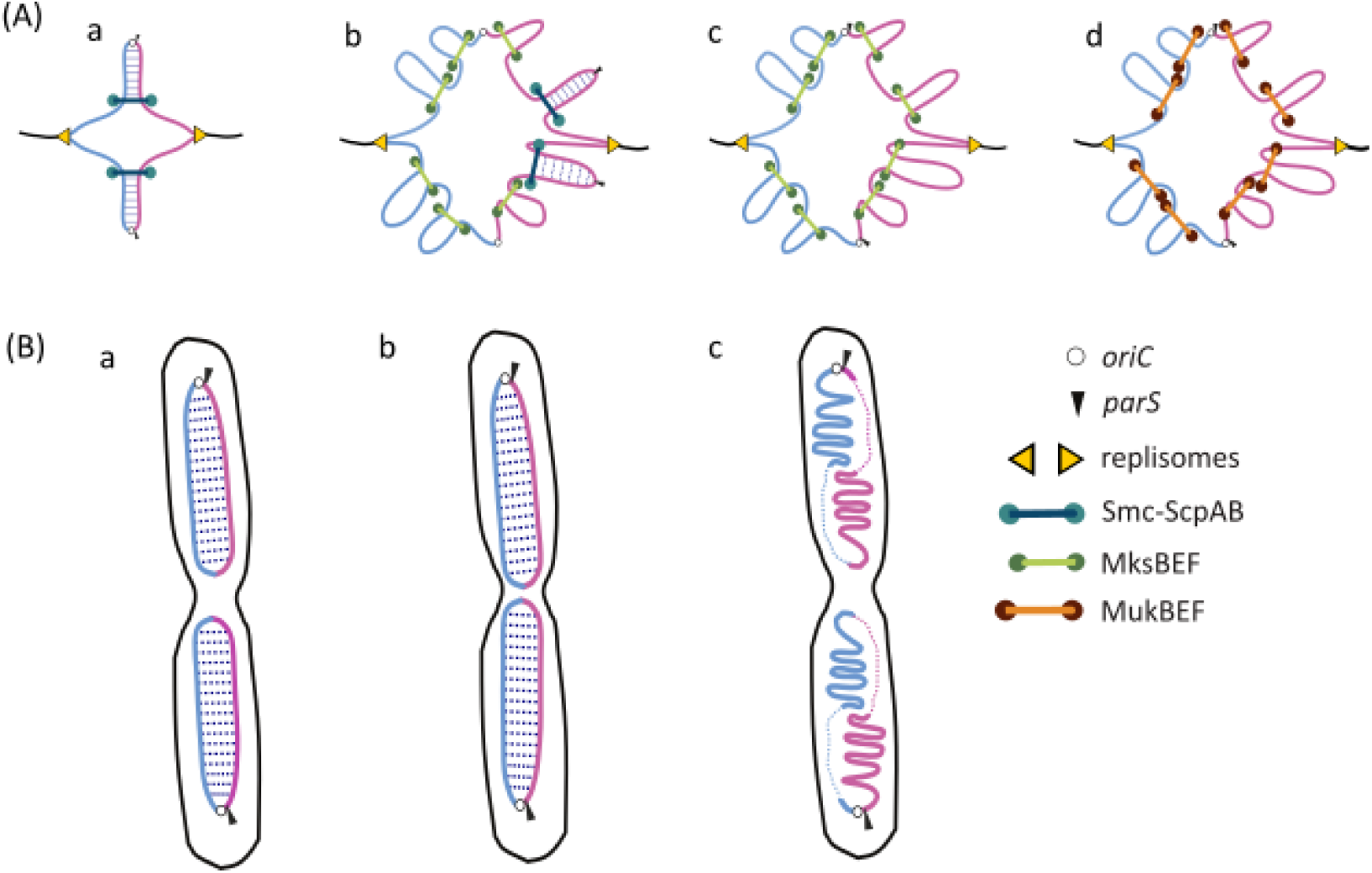
Model for bacterial condensin activity and interference in P. aeruginosa. **(A)** Schematized representation of replication arm alignment versus long-range contacts. The replication forks are represented by yellow arrowheads, oriC by a yellow circle, parS sites by a blue arrowhead duplicated DNA in red; Smc-ScpAB, MksBEF and MukBEF are depicted as blue, green and orange rings, respectively. (a) In a wt chromosome configuration, Smc-ScpAB translocation from *parS* sites promotes the juxtaposition of chromosome arms. It is not known how Smc-ScpAB prevents MksBEF long range contacts (see text). (b) When oriC is distant from *parS* sites, MksBEF creates long-range contacts in regions where it can act before Smc-ScpAB can align them. This implies (i) that long-range contacts caused by MksBEF occur between DNA replication and Smc-ScpAB action and (ii) that Smc-ScpAB acts from *parS* upon its replication. (c) In the absence of Smc-ScpAB, in a wt chromosome configuration, MksBEF can produce long-range contacts all over the chromosome, presumably in a progressive manner and following the replication process (see (b)). As MksBEF promotes the extension of the range of cis contacts in the absence of the juxtaposition of chromosome arms, a reduced processivity or stability of MksBEF onto DNA compared to Smc-ScpAB may generate multiple loops in the bacterial chromosome, i.e. an increase of long-range *cis* contacts. (d) In the presence of *E. coli* MukBEF, juxtaposition of chromosome arms by Smc-ScpAB is prevented and long-range contacts are generated by MukBEF all over the chromosome. **(B)** Schematized representation of chromosome segregation and disposition by bacterial condensins. (a) The ParABS system drives segregation of Ori regions to 20-80 % of the cell and translocation of Smc-ScpAB from *parS* sites promotes the juxtaposition of chromosome arms and a longitudinal organization of the chromosome. The position of *parS* site(s) determines the orientation of the chromosome in the cell. (b) In the absence of MksBEF, the longitudinal disposition of the chromosome is maintained and the absence of MksBEF affects the separation of loci in the Ter region of the chromosome. (c) In the absence of Smc-ScpAB, the juxtaposition of chromosome arms and the longitudinal disposition of the chromosome are lost with the two arms of the chromosome in separate halves of the cell.

### Alignment of chromosome arms by Smc-ScpAB controls chromosome disposition inside the cell

We demonstrate that in *P. aeruginosa*, Smc-ScpAB is critical for chromosome disposition inside the cell. We previously showed that a *parS* site localizes near the 0.2/0.8 relative cell length upon segregation by the ParAB complex (Vallet-Gely and Boccard, 2013), even when it is displaced from its native location and located farther away from *oriC* (Lagage et al., 2016). Here, we show that Smc-ScpAB initiates chromosome arm alignment from the displaced *parS* site, independently of its location on the genome. Therefore, changing the location of the *parS* site leads to a complete reorientation of the chromosome inside the cell, the region encompassing the *parS* site being localized near the 0.2/0.8 relative cell length and chromosome arm alignment by Smc-ScpAB being modified according to the modified *parS* location.

Remarkably, in the wild type genome configuration, inactivation of Smc-ScpAB dramatically affects the longitudinal arrangement of the chromosome: although the *parS* site remains localized near the 0.2/0.8 relative cell length, chromosome arms are no longer aligned along the long axis of the cell but rather occupy separate halves of the cell (Figure 2F). These results suggest that *P. aeruginosa* chromosome arms spontaneously arrange in opposite cell halves, and that Smc-ScpAB establishes a longitudinal organization by aligning chromosome arms (Figure 6B).

### MksBEF and MukBEF affect the organization of the bacterial chromosome in similar ways

Both MksBEF and MukBEF can extend the median range of *cis* contacts of *P. aeruginosa* chromosomal loci from 600 kb (in the wild type strain) to more than 850 kb (hereafter, we will refer to this extension as “long-range *cis* contacts formation”). To determine whether this range of *cis* contacts was specific to the *P. aeruginosa* chromosome or not, we applied the quantification method described here to data obtained in (Lioy et al., 2018) and measured the range of *cis* contacts in the presence or in the absence of MukBEF in *E. coli* (Figure S4E to S4G). It exceeds 900 kb when MukBEF is active and is reduced to 600 kb in its absence, indicating that very similar values are obtained for both bacteria.

These results are in agreement with the previous observation that MukBEF activity leads to an increase in the amount of contacts above 280 kb, and to the separation of the *E. coli* chromosome into two structurally distinct entities: one where MukBEF can act (outside of Ter), and one where MukBEF is prevented from acting by MatP (inside the Ter) (Lioy et al., 2018). Interestingly, our results in *P. aeruginosa* suggest that MukBEF is able to promote a similar higher-order chromosome organization in both organisms. This seems also to be the case for MksBEF, even though it is usually not apparent due to chromosome arm alignment by Smc-ScpAB. Therefore, our results indicate that the distant bacterial condensins MksBEF and MukBEF can generate similar higher-order chromosome organization in bacterial chromosomes.

The mechanism of DNA loop extrusion by eukaryotic SMC complexes was initially proposed twenty years ago (Nasmyth, 2001), and first modeled ten years later (Alipour and Marko, 2012). Since then, a considerable amount of evidence has accumulated, including direct visualization of loop formation *in vitro* by yeast condensin (Ganji et al., 2018) and human cohesin (Kim et al., 2019). Some questions remain concerning the mechanisms of DNA loop extrusion: for instance, whether loop are extruded asymmetrically or symmetrically, whether SMC complexes act as individual or higher order complexes, and whether or not loop extrusion requires external motors (for review, see (Hassler et al., 2018)). In bacteria, it has been proposed that Smc-ScpAB dependent arm alignment may also proceed by loop extrusion from its loading site to the chromosome (i.e. the *parS* sites) (Tran et al., 2017; Wang et al., 2017). It is tempting to speculate that MukBEF and MksBEF also proceed by DNA loop extrusion. Properties of these complexes may account for the different outcomes observed: an increase of the range of *cis* contacts compared to the alignment of replication arms by Smc-ScpAB. First, contrary to Smc-ScpAB, no specific loading site has been identified for MukBEF and MksBEF. These complexes might load onto the chromosome at different locations and extrude DNA, promoting long-range *cis* contact formation all over the chromosome (Figure 6A). Second, as observed for Condensin I that presents a shorter halftime on chromatin than Condensin II (Gerlich et al., 2006), a reduced processivity or stability of MukBEF and MksBEF onto DNA compared to Smc-ScpAB may generate multiple loops in the bacterial chromosome, i.e. an increase of long range *cis* contacts, as observed in metazoan mitotic chromosomes (Gibcus et al., 2018).

### Functional hierarchy between bacterial condensin activities

Factors modulating the activity of SMC complexes have already been described; for example, CTCF and cohesin cooperate to form chromatin loops and boundaries between TADs, while MatP prevents MukBEF activity in *E. coli*. Yet the molecular mechanisms at work remain to be characterized. In this study, we highlight interferences between bacterial SMC complexes, the first one between Smc-ScpAB and MksBEF (MksBEF can extend the range of *cis* contacts of chromosomal loci only when they are not aligned by Smc-ScpAB), and the second between MukBEF and Smc-ScpAB (MukBEF prevents Smc-ScpAB from aligning chromosome arms). Assuming that all three bacterial condensins work by loop extrusion, two hypotheses might explain the hierarchy in their activities: either the DNA conformation produced by one is not a good substrate for the other; or there is a direct competition between the different complexes acting on DNA. Considering that the range of *cis* contacts promoted by MukBEF and MksBEF are similar, and that Smc-ScpAB is able to align chromosomal loci presenting an extended range of *cis* contacts by MksBEF (in strains with rearranged chromosome configurations), it seems unlikely that the chromosome structuring generated by MukBEF prevents Smc-ScpAB from aligning chromosome arms. In contrast, we observed contacts between approximatively 300 kb on both sides of *oriC* on the contact map of the strain in which the *mukFEB* genes replaced the *mksFEB* genes, but not on the contact map of the Δ*smc* mutant with the same substitution (Figure 5E and 5F). This suggests that Smc-ScpAB might be able to initiate chromosome arm alignment in the presence of MukBEF, but not to progress far away from its loading site. Therefore, we favor a competition between the two complexes to explain the hierarchy observed between MukBEF and Smc-ScpAB upon introduction of the former in *P. aeruginosa.* Concerning MksBEF and Smc-ScpAB interference, it is tempting to speculate that in a wt chromosome configuration, the early engagement of Smc-ScpAB at *parS* sites and its subsequent association with newly replicated DNA might prevent MksBEF to act on newly replicated DNA and produce long-range contacts. Alternatively, the juxtaposed replication arms established by Smc-ScpAB might not be a good substrate for MksBEF (Figure 6A).

### Distinct activities of bacterial condensins in chromosome segregation

Here we show that both Smc-ScpAB and MksBEF contribute to the separation of distal loci in the presence of the ParABS system. However, in the absence of the ParABS system, the two condensins are not equivalent as MksBEF allows a higher growth rate and a low level of anucleate cells compared to Smc-ScpAB (4% versus 25% anucleate cells). Strikingly, MukBEF can substitute for MksBEF in *P. aeruginosa*. Molecular mechanisms responsible for chromosome segregation by MukBEF and MksBEF remain to be characterized. Considering that both complexes promote formation of long-range *cis* contacts, it is tempting to speculate that this is how they contribute to segregation. However, we did not detect extension of the range of *cis* contacts by MksBEF when chromosome arms are aligned by Smc-ScpAB, for example in the Δ*parA* mutant in which MksBEF is critical for chromosome segregation. This indicates either that long-range *cis* contact formation by MksBEF is not involved in chromosome segregation, or that MksBEF activity required for chromosome segregation, involving either a low level of long-range contacts over the chromosome or contacts occurring at a few loci, cannot be detected using the 3C-approach.

In agreement with the observation that most bacteria in which MukBEF has replaced Smc-ScpAB lack a ParABS system, MukBEF is thought to be able to sustain correct chromosome segregation by itself, as in *E. coli* for instance. Remarkably, the efficiency for chromosome segregation of MukBEF in *P. aeruginosa* increases with the growth rate. These observations lead us to the hypothesis that DNA topology or chromatin organization in *P. aeruginosa* might not be optimal for MukBEF activity. Considering that differences in the level of supercoiling have been proposed to explain differences in phenotypes associated with MukBEF in *E. coli* and *Salmonella* (Higgins, 2016), it is tempting to speculate that the failure of MukBEF or MksBEF to sustain optimal chromosome segregation in *P. aeruginosa* roots from a different state of DNA topology.

Considering that MksBEF is present in all *Pseudomonas* species, it is likely to be critical for *Pseudomonas* survival in the diverse environments that these ubiquitous bacteria colonize. It might constitute a useful backup of the ParABS system in environments in which this system might not be optimal for chromosome segregation. Our results indicate that the separation of markers in the terminal region of the chromosome is affected in the absence of MksBEF whereas the ParABS system is most efficient for Ori region separation and positioning (Figure 6B). This would imply that *Pseudomonas* species have evolved with two major bacterial chromosome segregation systems, one specifically driving segregation of the Ori region and the other promoting separation of distal bulk chromosome.

## Acknowledgments

We thank Stéphane Duigou, Jean-Luc Ferat and Yoshiharu Yamaichi for careful reading of the manuscript; members of the FB laboratory for fruitful discussions. We also thank Valentin V. Rybenkov for sharing unpublished results, Christine Pourcel for providing the PAO1_orsay strain and Stéphane Duigou for the gift of the pBAD::*mukFEB* plasmid. We thank the I2BC genomic facility for high-throughput sequencing. We thank François-Xavier Barre for sharing Matlab functions. This research was supported by CNRS and Université Paris-Sud.

## Author Contributions

Conceptualization, V.S.L., F.B. and I.V.; Methodology, V.S.L. and I.V.; Investigation, V.S.L., V.L. and I.V; Resources, I. J.; Writing - Draft, V.S.L., F.B. and I.V.; Writing – Reviewing and Editing, V.S.L., F.B. and I.V.; Funding acquisition, F.B.; Supervision, F.B and I.V.

## Materials and Methods

### Bacterial Strains and plasmids

The PAO1 strain that we used in our previous studies was provided by Arne Rietsch (Case Western Reserve University). As described in (Lagage et al., 2016), this PAO1 strain does not present the inversion described for the sequenced PAO1-UW subclone resulting from homologous recombination between the *rrnA* and *rrnB* loci, which are orientated in opposite directions and separated by 2.2 Mbp (Stover et al, 2004). It also contains the 12 kb insertion and 1006 bp deletion described in (Klockgether et al., 2010). Considering that the 1006 bp deletion encompassed *mksE* and *mksF*, two genes that were potentially important for chromosome maintenance, we decided to reintroduce them in our PAO1 strain (see below) to obtain a wild type strain background containing the three systems potentially important for segregation, the ParABS system and the two bacterial condensins MksBEF and Smc-ScpAB.

*Escherichia coli* DH5α (Invitrogen) was used as the recipient strain for all plasmid constructions, whereas *E. coli* strain β2163 (Demarre et al., 2005) was used to mate plasmids into *P. aeruginosa*. All the integration vectors carry the mobilization region from RP4, the ColE1 origin of replication and the *aacC1* gene (conferring resistance to gentamicin). Plasmids derived from pEXG2 (Rietsch et al., 2005) also contain the *sacB* gene for allelic exchange.

Plasmids allowing insertion of chromosomal tags and their visualization using fluorescent proteins have been described previously (Vallet-Gely and Boccard, 2013).

The deletion construct for *smc* was generated by amplifying flanking regions by the PCR and then splicing the flanking regions together by overlap extension PCR, replacing the *smc* gene by a 6-bp linker sequence 5’-GAATTC-3’. The resulting PCR products were cloned on *Xba*I/*Hind*III fragments into plasmid pEXG2, yielding plasmid pEXMΔ*smc*. This plasmid was then used to create strains Δ*mks* Δ*smc*, Δ*parS* Δ*smc, parS* ^+550^ Δ*smc, parS* ^-330^ Δ*smc* and *oriC ins1* Δ*smc* by allelic exchange. Deletions were confirmed by PCR. The deletion construct for the *parA* was generated by amplifying flanking regions by PCR and then splicing the flanking regions together by overlap extension PCR, replacing the *parA* gene by a 6-bp linker sequence 5’-GAATTC-3’. The resulting PCR products were cloned on *Xba*I/*Hind*III fragments into plasmid pEXG2, yielding plasmid pEXMΔ*parA*. This plasmid was then used to create strains Δ*parA* by allelic exchange. Deletion was confirmed by PCR. The insertion construct for the ectopic *oriC* was generated by amplifying the two flanking regions of the insertion site (called *ins1*) by PCR and then splicing the flanking regions together by overlap extension PCR, introducing an *EcoR*I restriction site at the insertion site. The resulting PCR product was cloned on a *Xba*I/*Hind*III fragment into plasmid pEXG2, yielding plasmid pEXM*ins1*. A 600 pb fragment encompassing *oriC* was amplified using PCR and cloned on an *EcoR*I fragment into plasmid pEXM*ins1*, yielding plasmid pEXM*oriCins1*. This plasmid was then used to create strains *oriC ins1* Δ*mks* by allelic exchange. Insertion was confirmed by PCR. The insertion construct for the *mksEF* genes was generated by amplifying the 1006 pb deletion and flanking regions from the PAO1_Orsay strain (Latino et al., 2016) by PCR. The resulting PCR product was cloned on a *Hind*III/*Xba*I fragment into plasmid pEXG2, yielding plasmid pEXMksEF^IN^. This plasmid was then used to create strains WT, Δ*smc*, Δ*parS*, Δ *parS123, parS* ^+550^, *parS* ^-330^ and *oriC ins1*. Insertions were confirmed by PCR. The construct used for replacing the *mksFEB* genes of *P. aeruginosa* by the *mukFEB* genes of *E. coli* (from the ATG of *mksF*/*mukF* to the TGA of *mksB*/*mukB*) was generated as follow. Overlapping PCR was used to splice together the region upstream of *mksF* in *P. aeruginosa* and the beginning of the *mukF* gene of *E. coli* on the one hand, and the end of the *mukB* gene of *E. coli* together with the downstream region of *mksB* in *P. aeruginosa* on the other hand. The two resulting fragments were cloned together in the pEXG2 plasmid on a *Xba*I/*Pac*I fragment, separated by an *EcoR*I site, yielding the pEXpreMuk plasmid. The rest of the mukFEB operon was cloned as a SacI/KpnI fragement in the pEXpreMuk plasmid, yielding the pEXMukFEB^IN^ plasmid. This plasmid was then used to create strains Δ*mks::mukFEB*, Δ*smc* Δ*mks::mukFEB, parS*^+550^ Δ*mks::mukFEB* and *oriC ins1* Δ*mks::mukFEB*. Insertions were confirmed by PCR.

All strains used in this study are listed in the Supplementary Table 1.

### Media and growth conditions

Unless otherwise stated, all experiments were performed in minimal medium A supplemented with 0.25% citrate as carbone source. In this growth medium, wild type cells contain a single replicating chromosome: replication just started in newborn cells, whereas division occurred shortly after completion of the replication process. For fluorescent microscopy experiments, strains were grown overnight in LB, diluted 300 times in minimal medium A supplemented with 0.25% citrate and grown at 30°C until they reached an OD600 comprised between 0.05 and 0.1. IPTG was added to growth medium at 0.5 mM for observation of chromosomal tags. Observation of chromosomal loci was performed as described previously (Lagage et al., 2016). For chromosome capture experiments, strains were grown overnight in LB, diluted 500 times in minimal medium A supplemented with 0.25% citrate and grown at 30°C until they reach an OD600 of 0.1.

For anucleate cell quantification, after overnight cultures in LB, cells were diluted 300 time and grown in different growth medium (LB, minimal medium A supplemented with 0.2% glucose and 0.12% of casamino acids, or minimal medium A supplemented with 0.25% citrate) until OD600 0.1, fixed with an equal volume of a 1×PBS solution containing 5% paraformaldehyde and 0.06% glutaraldehyde, and processed as described previously (Lagage et al., 2016).

### Chromosome conformation capture

3C libraries were generated as described previously (Lioy and Boccard 2018). Briefly, 100 ml of culture was crosslinked with fresh formaldehyde for 30 minutes (5% final concentration) at room temperature (RT) followed by 30 minutes at 4°C. Formaldehyde was quenched with a final concentration of 0.25 M glycine for 20 minutes at 4°C. Fixed cells were collected by centrifugation, frozen on dry ice and stored at -80°C until use. Frozen pellets of ≈ 1-2 × 10^9^ cells were thawed, suspended in 600 µl Tris 10 mM EDTA 0.5 mM (TE) (pH 8) with 4 µl of lyzozyme (35 U/µl; Tebu Bio), and incubated at RT for 20 minutes. SDS was added to the mix (final concentration 0.5%) and the cells incubated for 10 minutes at RT. 50µl of lysed cells were transferred to a tube containing 450µL of digestion mix (1X NEB 1 buffer, 1% triton X-100). This process was repeated 11 times (total 12 tubes with 500 µl). 100 units of HpaII were added to 10 tubes. All the tubes were then incubated for 2 hours at 37°C. To stop the digestion reaction, 8 tubes were immediately centrifuged during 20 min at 20,000 g, and pellets were suspended in 500µl of sterile water. The digested DNA (4 ml in total) was split in 4 aliquots, and diluted in 8 ml ligation buffer (1X ligation buffer NEB without ATP, 1 mM ATP, 0.1 mg/ml BSA, 125 Units of T4 DNA ligase 5 U/µl). Ligation was performed at 16°C for 4 hours, followed by incubation overnight (ON) at 65°C with 100 µl of proteinase K (20 mg/ml) and 100µl EDTA 500 mM. DNA was then precipitated with an equal volume of 3 M Na-Acetate (pH 5.2) and two volumes of isopropanol. After one hour at -80°C, DNA was pelleted, suspended in 500µl 1X TE buffer. The remaining 4 tubes (2 tubes with HpaII and 2 tubes without restriction enzyme) were directly incubated with 50 µl of proteinase K (20mg/ml) overnight at 65°C.Finally, all the tubes were transferred into 2 ml centrifuge tubes, extracted twice with 400 µl phenol-chloroform pH 8.0, precipitated, washed with 1 ml cold ethanol 70% and diluted in 30 µl 1X TE buffer in presence of RNAse A (1 μg/ml). Tubes containing the ligated DNA (3C libraries), the digested DNA or the non-digested DNA were pooled into 3 different tubes and the efficiency of the 3C preparation was assayed by running a 1% agarose gel. 3C libraries were quantified on the gel using QuantityOne software (BioRad).

### Processing of libraries for Illumina sequencing

Approximately 5 µg of a 3C library was suspended in water (final volume 130 µL) and sheared using a Covaris S220 instrument (Duty cycle 5, Intensity 5, cycles/burst 200, time 60 sec for 4 cycles). The DNA was purified using Qiaquick^®^ PCR purification kit, DNA ends were prepared for adapter ligation following standard protocols (see Cournac 2016). Custom-made adapters (Marbouty 2015) were ligated overnight at 4°C. Ligase was inactivated by incubating the tubes at 65°C for 20 minutes. To purify DNA fragments ranging in size from 400 to 900 pb, a PippinPrep apparatus (SAGE Science) was used. For each library, one PCR reaction of 12 cycles was performed (using 2 µL of 3C library, 0.2 µM Illumina primers PE1.0 and PE2.0 and 1 unit of Taq Phusion [Finnzymes]). The PCR product was purified on Qiagen MinElute columns and primers dimers were removed from the 3C library by using AMPure XP beads following the manufacturer’s protocol (Beckman Coulter). Finally, libraries were subjected to paired-end sequencing on an Illumina sequencer (NextSeq500).

### Processing of sequencing data

PCR duplicates from each 3C library sequence dataset were discarded using the 6 Ns of custom-made adapters (Marbouty 2015). Reads were aligned independently using Bowtie 2 in very sensitive mode (Langmead and Salzberg, 2012). Only reads with mapping quality > 30 were kept to establish contact maps.

### Marker Frequency Analysis

MFA was performed on gDNAs extracted from cells grown in the same conditions as for the chromosome capture experiments, using the Sigma GenElute^®^ bacterial genomic DNA kit. Libraries for sequencing were prepared following Illumina TruSeq protocol (Westburg). Libraries were sequenced on an Illumina NextSeq500 instrument, following manufacturer’s protocol. Approximatively 15 million of reads were recovered for each sample. Reads can be downloaded using the GEO number XXXX. We used the Bowtie2 software to perform the mapping in the local mode, and the mpileup software from Samtools to calculate the coverage for each genome position. Then, as described in Galli et al., 2019, enrichment of uniquely mapping sequence reads were calculated over 1kb and 200kb sliding windows. Local 1kb-window values deviating by more than 15% from the local 200kb-window values were discarded from the analysis.

### Generation of contact maps

Contact maps were built as describe previously (Lioy et al., 2018). Briefly, each read was assigned to a restriction fragment. Non-informative events such as self-circularized restriction fragments, or uncut co-linear restriction fragments were discarded (Cournac et al., 2012). The genome was then binned into 10 kb units and the corresponding contact map generated and normalized through the sequential component normalization procedure (SCN; Cournac et al., 2012). Contact maps are visualized as log matrices, to facilitate visualization.

### Quantification of the range of *cis* contacts along the arms

We used a three steps process based on the one described in Wang et al., 2017 to determine the endpoint of DNA juxtaposition on Hi-C maps, although we used it to determine the width of the primary diagonal. First, we calculated the median of the contact map, and to estimate the standard deviation (σ) we used a robust statistic which says that σ = 1.4826*mad (mad = median absolute deviation) (Rousseeuw and Croux, 1993). For each contact map, thresholds equal to 0.1, 0.2, 0.25, 0.5 and 0.75 times the standard deviation (σ) above the median were set. Contact frequencies above or below the threshold were assigned a value of 1 or 0, respectively, generating a binary contact map (see Sup. Figure 3). Next, we used a point connecting algorithm, in order to be able to discriminate background from significant interactions. The size of each connected element identified by the “bwlabel” function of matlab was calculated. Most of them are very small (around 98% contain less than 10 points). We fixed another arbitrary limit of 30 points in a connected element to be considered as significant (see Sup. Figure 3). Finally, mainly to facilitate the measuring process, we used the “imclose()” function of matlab to fill out the empty points comprised in the connected elements, using a diamond shape with a size of 5.

Then, the width of the primary diagonal was calculated for each point of a chromosome arm that is not located in the Ter or Ori regions (where *cis* contacts are obviously extended by the encounter between the two diagonals). For comparison purposes, we fixed the limits of the chromosomal regions according to the chromosome configuration. For the wild type configuration (strains WT, Δ*mks*, Δ*smc*, Δ*parS*, Δ*smc* Δ*parS*, Δ*smc* Δ*mks*, Δ*smc* Δ*mks::mukBEF*, Δ*mks::mukBEF* and Δ*parA*), loci located from - 1.25 to 2.25 Mb on the left of *oriC* constitute the “left arm”, whereas loci located from 1.2 to 2.1 Mb on the right of *oriC* constitute the “right arm”. For the *parS* ^+550^ configuration (strains *parS* ^+550^, *parS* ^+550^ Δ*mks, parS* ^+550^ Δ*smc* and *parS* ^+550^ Δ*mks::mukFEB*), loci located from 0.9 to 2.8 Mb on the left of *oriC* constitute the “left arm”, whereas loci located from 1.5 to 2.4 Mb on the right of *oriC* constitute the “right arm”. For the *parS* ^-330^ configuration (strains *parS* ^-330^, *parS* ^-330^ Δ*mks* and *parS* ^-330^ Δ*smc*), loci located from 1.1 to 2.4 Mb on the left of *oriC* constitute the “left arm”, whereas loci located from 0.9 to 1.9 Mb on the right of *oriC* constitute the “right arm”. For the *oriC ins1* configuration (strains *oriC ins1, oriC ins1* Δ*mks* and *oriC ins1* Δ*smc*), loci located from 1.1 to 2.1 Mb on the left of *oriC* constitute the “left arm”, whereas loci located from 1.2 to 2.3 Mb on the right of *oriC* constitute the “right arm”.

The range of cis contact was estimated from the width of the primary diagonal by multiplying the number of measured bins by the size of the bin (10 kb), and dividing by two (this range is symmetric on both sides of the chromosomal locus considered). A boxplot representation is then used, allowing visualization of the whole range of *cis* contacts for every considered chromosome region. Remarkably, despite the fact that replicate contact maps may present different level of “background noise”, this method to estimate the range of *cis* contacts along the chromosome arms gives similar results.

We chose the threshold of 0.2*σ for the main figures, as it fitted best with the observed contact maps. However, considering the arbitrary nature of this threshold, we also analyzed all contact maps using the 0.1, 0.25 and 0.5 thresholds. The entirety of these results are presented in Figure S4A, B, C and D, to facilitate comparisons. They show that the estimated range of *cis* contacts along the arms differs from one threshold to another; however, the differences observed between the different mutants are mostly conserved, independently of the chosen threshold.

To further validate this approach, we also used it to quantify the range of *cis* contacts along the arms of the *E. coli* chromosome, in the Ter region and elsewhere, in the wild type strain, the Δ*matP* mutant and the Δ*mukB* mutant. Results using 0.2, 0.25 and 0.5 as the first threshold are presented Figure S4E, F and G respectively. They confirm that MukBEF is responsible for long range contacts outside the Ter region, and that MatP prevents the formation of these long range contacts in the Ter region (Lioy et al., 2018). Here also, the threshold affects the extent of *cis* contacts, but not the effect of the absence of MatP and MukBEF.

## DATA AND SOFTWARE AVAILABILITY

Contact maps and FASTQ files of the reads were deposited in the NCBI database under the GEO accession number ### (pending).

## References

Alipour, E., and Marko, J.F. (2012). Self-organization of domain structures by DNA-loop-extruding enzymes. Nucleic Acids Res. 40, 11202–11212.

Badrinarayanan, A., Le, T.B., and Laub, M.T. (2015). Bacterial chromosome organization and segregation. Annu Rev Cell Dev Biol 31, 171–199.

Battistuzzi, F.U., Feijao, A., and Hedges, S.B. (2004). A genomic timescale of prokaryote evolution: insights into the origin of methanogenesis, phototrophy, and the colonization of land. BMC Evol. Biol. 4, 44.

Böhm, K., Giacomelli, G., Schmidt, A., Imhof, A., Koszul, R., Marbouty, M., and Bramkamp, M. (2020). Chromosome organization by a conserved condensin-ParB system in the actinobacterium Corynebacterium glutamicum. Nat Commun 11, 1485.

Brézellec, P., Hoebeke, M., Hiet, M.-S., Pasek, S., and Ferat, J.-L. (2006). DomainSieve: a protein domain-based screen that led to the identification of dam-associated genes with potential link to DNA maintenance. Bioinformatics 22, 1935–1941.

Cobbe, N., and Heck, M.M.S. (2004). The evolution of SMC proteins: phylogenetic analysis and structural implications. Mol. Biol. Evol. 21, 332–347.

Cournac, A., Marie-Nelly, H., Marbouty, M., Koszul, R., and Mozziconacci, J. (2012). Normalization of a chromosomal contact map. BMC Genomics 13, 436.

Cournac, A., Marbouty, M., Mozziconacci, J., and Koszul, R. (2016). Generation and Analysis of Chromosomal Contact Maps of Yeast Species. In Yeast Functional Genomics, F. Devaux, ed. (Springer New York), pp. 227–245.

Doron, S., Melamed, S., Ofir, G., Leavitt, A., Lopatina, A., Keren, M., Amitai, G., and Sorek, R. (2018). Systematic discovery of antiphage defense systems in the microbial pangenome. Science 359.

Ganji, M., Shaltiel, I.A., Bisht, S., Kim, E., Kalichava, A., Haering, C.H., and Dekker, C. (2018). Real-time imaging of DNA loop extrusion by condensin. Science 360, 102–105.

Gibcus, J.H., Samejima, K., Goloborodko, A., Samejima, I., Naumova, N., Nuebler, J., Kanemaki, M.T., Xie, L., Paulson, J.R., Earnshaw, W.C., et al. (2018). A pathway for mitotic chromosome formation. Science 359.

Gruber, S., and Errington, J. (2009). Recruitment of condensin to replication origin regions by ParB/SpoOJ promotes chromosome segregation in B. subtilis. Cell 137, 685–696.

Hassler, M., Shaltiel, I.A., and Haering, C.H. (2018). Towards a Unified Model of SMC Complex Function. Curr. Biol. 28, R1266–R1281.

Higgins, N.P. (2016). Species-specific supercoil dynamics of the bacterial nucleoid. Biophys Rev 8, 113–121.

Jalal, A.S., Tran, N.T., and Le, T.B. (2020). ParB spreading on DNA requires cytidine triphosphate in vitro. Elife 9.

Kawalek, A., Wawrzyniak, P., Bartosik, A.A., and Jagura-Burdzy, G. (2020). Rules and Exceptions: The Role of Chromosomal ParB in DNA Segregation and Other Cellular Processes. Microorganisms 8.

Kim, Y., Shi, Z., Zhang, H., Finkelstein, I.J., and Yu, H. (2019). Human cohesin compacts DNA by loop extrusion. Science 366, 1345–1349.

Kleckner, N., Fisher, J.K., Stouf, M., White, M.A., Bates, D., and Witz, G. (2014). The bacterial nucleoid: nature, dynamics and sister segregation. Curr. Opin. Microbiol. 22, 127–137.

Lagage, V., Boccard, F., and Vallet-Gely, I. (2016). Regional Control of Chromosome Segregation in Pseudomonas aeruginosa. PLoS Genet. 12, e1006428.

Langmead, B., and Salzberg, S.L. (2012). Fast gapped-read alignment with Bowtie 2. Nat Methods 9, 357–359.

Le, T.B.K., Imakaev, M.V., Mirny, L.A., and Laub, M.T. (2013). High-resolution mapping of the spatial organization of a bacterial chromosome. Science 342, 731–734.

Lioy, V.S., Cournac, A., Marbouty, M., Duigou, S., Mozziconacci, J., Espéli, O., Boccard, F., and Koszul, R. (2018). Multiscale Structuring of the E. coli Chromosome by Nucleoid-Associated and Condensin Proteins. Cell 172, 771-783.e18.

Livny, J., Yamaichi, Y., and Waldor, M.K. (2007). Distribution of centromere-like parS sites in bacteria: insights from comparative genomics. J. Bacteriol. 189, 8693–8703.

Mäkelä, J., and Sherratt, D.J. (2020). Organization of the Escherichia coli Chromosome by a MukBEF Axial Core. Mol. Cell.

Marbouty, M., Le Gall, A., Cattoni, D.I., Cournac, A., Koh, A., Fiche, J.-B., Mozziconacci, J., Murray, H., Koszul, R., and Nollmann, M. (2015). Condensin- and Replication-Mediated Bacterial Chromosome Folding and Origin Condensation Revealed by Hi-C and Super-resolution Imaging. Mol. Cell 59, 588–602.

Mercier, R., Petit, M.-A., Schbath, S., Robin, S., El Karoui, M., Boccard, F., and Espéli, O. (2008). The MatP/matS site-specific system organizes the terminus region of the E. coli chromosome into a macrodomain. Cell 135, 475–485.

Minnen, A., Bürmann, F., Wilhelm, L., Anchimiuk, A., Diebold-Durand, M.-L., and Gruber, S. (2016). Control of Smc Coiled Coil Architecture by the ATPase Heads Facilitates Targeting to Chromosomal ParB/parS and Release onto Flanking DNA. Cell Rep 14, 2003–2016.

Nasmyth, K. (2001). Disseminating the genome: joining, resolving, and separating sister chromatids during mitosis and meiosis. Annu. Rev. Genet. 35, 673–745.

Niki, H., Jaffé, A., Imamura, R., Ogura, T., and Hiraga, S. (1991). The new gene mukB codes for a 177 kd protein with coiled-coil domains involved in chromosome partitioning of E. coli. EMBO J 10, 183–193.

Nolivos, S., and Sherratt, D. (2014). The bacterial chromosome: architecture and action of bacterial SMC and SMC-like complexes. FEMS Microbiol Rev 38, 380–392.

Nolivos, S., Upton, A.L., Badrinarayanan, A., Müller, J., Zawadzka, K., Wiktor, J., Gill, A., Arciszewska, L., Nicolas, E., and Sherratt, D. (2016). MatP regulates the coordinated action of topoisomerase IV and MukBEF in chromosome segregation. Nat Commun 7, 10466.

Osorio-Valeriano, M., Altegoer, F., Steinchen, W., Urban, S., Liu, Y., Bange, G., and Thanbichler, M. (2019). ParB-type DNA Segregation Proteins Are CTP-Dependent Molecular Switches. Cell 179, 1512-1524.e15.

Panas, M.W., Jain, P., Yang, H., Mitra, S., Biswas, D., Wattam, A.R., Letvin, N.L., and Jacobs, W.R. (2014). Noncanonical SMC protein in Mycobacterium smegmatis restricts maintenance of Mycobacterium fortuitum plasmids. Proc. Natl. Acad. Sci. U.S.A. 111, 13264–13271.

Petrushenko, Z.M., She, W., and Rybenkov, V.V. (2011). A new family of bacterial condensins. Mol. Microbiol. 81, 881–896.

Sawitzke, J.A., and Austin, S. (2000). Suppression of chromosome segregation defects of Escherichia coli muk mutants by mutations in topoisomerase I. Proc. Natl. Acad. Sci. U.S.A. 97, 1671–1676.

Soh, Y.-M., Davidson, I.F., Zamuner, S., Basquin, J., Bock, F.P., Taschner, M., Veening, J.-W., De Los Rios, P., Peters, J.-M., and Gruber, S. (2019). Self-organization of parS centromeres by the ParB CTP hydrolase. Science 366, 1129–1133.

Sullivan, N.L., Marquis, K.A., and Rudner, D.Z. (2009). Recruitment of SMC by ParB-parS organizes the origin region and promotes efficient chromosome segregation. Cell 137, 697–707.

Tran, N.T., Laub, M.T., and Le, T.B.K. (2017). SMC Progressively Aligns Chromosomal Arms in Caulobacter crescentus but Is Antagonized by Convergent Transcription. Cell Rep 20, 2057–2071.

Uhlmann, F. (2016). SMC complexes: from DNA to chromosomes. Nat. Rev. Mol. Cell Biol. 17, 399–412.

Vallet-Gely, I., and Boccard, F. (2013). Chromosomal organization and segregation in Pseudomonas aeruginosa. PLoS Genet. 9, e1003492.

Wang, X., Montero Llopis, P., and Rudner, D.Z. (2013). Organization and segregation of bacterial chromosomes. Nat. Rev. Genet. 14, 191–203.

Wang, X., Le, T.B.K., Lajoie, B.R., Dekker, J., Laub, M.T., and Rudner, D.Z. (2015). Condensin promotes the juxtaposition of DNA flanking its loading site in Bacillus subtilis. Genes Dev. 29, 1661–1675.

Wang, X., Brandão, H.B., Le, T.B.K., Laub, M.T., and Rudner, D.Z. (2017). Bacillus subtilis SMC complexes juxtapose chromosome arms as they travel from origin to terminus. Science 355, 524–527.

Wang, X., Hughes, A.C., Brandão, H.B., Walker, B., Lierz, C., Cochran, J.C., Oakley, M.G., Kruse, A.C., and Rudner, D.Z. (2018). In Vivo Evidence for ATPase-Dependent DNA Translocation by the Bacillus subtilis SMC Condensin Complex. Mol. Cell 71, 841-847.e5.

Wilhelm, L., Bürmann, F., Minnen, A., Shin, H.-C., Toseland, C.P., Oh, B.-H., and Gruber, S. (2015). SMC condensin entraps chromosomal DNA by an ATP hydrolysis dependent loading mechanism in Bacillus subtilis. Elife 4.

