## Supplementary material for "Distinct activities of bacterial condensins for chromosome management in *Pseudomonas aeruginosa*": Supplemantal Table

**Supplementary Table 1**

| Experimental Models: Organisms/Strains | | |
|---|---|---|
| IVGB578 (PAO1 *mksEF^IN^)* | This paper | WT |
| IVGB579 (PAO1 Δ*smc mksEF^IN^)* | This paper | Δ*smc* |
| PAO1 | (Lagage et al., 2016) | Δ*mks* |
| IVGB464 (PAO1 Δ*smc)* | This paper | Δ*mks* Δ*smc* |
| IVGB580 (PAO1 Δ*parS mksEF^IN^)* | This paper | Δ*parS* |
| IVGB941(PAO1 Δ*parS123 mksEF^IN^)* | This paper | Δ*parS 123* |
| IVGB735 (PAO1Δ*parA mksEF^IN^* ) | This paper | Δ*parA* |
| IVGB655 **(**PAO1 Δ*parS mksEF^IN^* Δ*smc)* | This paper | Δ*parS* Δ*smc* |
| VLB1 **(**PAO1 Δ*parS)* | (Lagage et al., 2016) | Δ*parS* Δ*mks* |
| IVGB982 **(**PAO1 Δ*mks::mukFEB)* | This paper | Δ*mks::mukFEB* |
| IVGB984 **(**PAO1 Δ*parS* Δ*mks::mukFEB)* | This paper | Δ*parS* Δ*mks::mukFEB* |
| IVGB988 **(**PAO1 Δ*mks::mukFEB* (IVGB982) Δ*smc)* | This paper | Δ*smc* Δ*mks::mukFEB* |
| IVGB717(PAO1 Δ*parS parS^+550^ mksEF^IN^)* | This paper | *parS* ^+550^ |
| IVGB480 **(**PAO1 Δ*parS parS^+550^*) | (Lagage et al., 2016) | *parS^+550^* Δ*mks* |
| IVGB990(PAO1 Δ*parS parS^+550^ mksEF^IN^* Δ*smc*) | This paper | *parS^+550^* Δ*smc* |
| IVGB994(PAO1 Δ*parS parS^+550^* Δ*mks::mukFEB)* | This paper | *parS^+550^* Δ*mks::mukFEB* |
| IVGB942(Δ*parS parS ^-330^* Δ*rrnD mksEF^IN^)* | This paper | *parS^-330^* |
| IVGB524(Δ*parS parS ^-330^* Δ*rrnD)* | (Lagage et al., 2016) | *parS^-330^* Δ*mks* |
| IVGB991(PAO1 *oriC ins1* (IVGB398) *mksEF^IN^)* | This paper | *parS^-330^* Δ*smc* |
| IVGB895(PAO1 *oriC ins1* (IVGB398) *mksEF^IN^)* | This paper | *oriC ins1* |
| IVGB398(PAO1 *oriC ins1)* | This paper | *oriC ins1* Δ*mks* |
| IVGB1003(PAO1 *oriC ins1 mksEF^IN^* Δ*smc)* | This paper | *oriC ins1* Δ*smc* |

| | | |
|---|---|---|
| IVGB1013(PAO1 *oriC ins1 Δmks::mukFEB)* | This paper | *oriC ins1*<br>*Δmks::mukFEB* |
| IVGB1022 [PAO1 *oriC ins1* parST1-PA5480(92-L) tetO-PA4457(1,275-R)] | This paper | *oriC ins1 Δmks*<br>92-L & 1,275-L |
| IVGB1023 [PAO1 *oriC ins1 mksEF$^{IN}$* parST1-PA5480(92-L) tetO-PA4457(1,275-R)] | This paper | *oriC ins1*<br>92-L & 1,275-L |
| IVGB1024 [PAO1 *oriC ins1 Δmks::mukFEB* parST1-PA5480(92-L) tetO-PA4457(1,275-R)] | This paper | *oriC ins1*<br>*Δmks::mukFEB*<br>92-L & 1,275-L |
| IVGB1025 [PAO1 *oriC ins1 Δmks::mukFEB* parST1-PA5480(92-L) tetO-PA4027(1,006-R)] | This paper | *oriC ins1*<br>*Δmks::mukFEB*<br>92-L & 1,006-R |
| IVGB1026 [PAO1 *Δmks::mukFEB* parST1-PA5480(92-L) tetO-PA4457(1,275-R)] | This paper | *Δmks::mukFEB*<br>92-L & 1,275-L |
| IVGB1027 [PAO1 *Δmks::mukFEB* parST1-PA5480(92-L) tetO-PA4027(1,006-R)] | This paper | *Δmks::mukFEB*<br>92-L & 1,006-R |
| IVGB119 [PAO1 parST1-PA3573(1,509-R) tetO-PA4457(1,275-L)] | (Vallet-Gely and Boccard, 2013) | *Δmks*<br>1,275-L & 1,509-R |
| IVGB120 [PAO1 parST1-PA5480(92-L) tetO-PA0069(82-R)] | (Vallet-Gely and Boccard, 2013) | *Δmks*<br>3,090-L & 2,857-R |
| IVGB122 [PAO1 parST1-PA5480(92-L) tetO-PA0069(82-R)] | This paper | *Δmks*<br>82-R & 92-L |
| IVGB125 [PAO1 parST1-PA3573(1,509-R) tetO-PA0981(1,812-L)] | (Vallet and Boccard, 2013) | *Δmks*<br>1,509-R & 1,812-L |
| IVGB127 [PAO1 parST1-PA0981(1,812-L) tetO-PA4457(1,275-L)] | (Vallet-Gely and Boccard, 2013) | *Δmks*<br>1,275-L & 1,812-L |
| IVGB170 [PAO1 parST1-PA0572(628-R) tetO-PA4027(1,006-R)] | (Vallet-Gely and Boccard, 2013) | *Δmks*<br>1,006-R |
| IVGB174 [PAO1 parST1-PA4457(1,275-L) tetO-PA4822(851-L)] | (Vallet-Gely and Boccard, 2013) | *Δmks*<br>1,275-L |
| IVGB247 [PAO1 parST1-PA1428(2,302-L) tetO-PA0981(1,812-L)] | (Vallet-Gely and Boccard, 2013) | *Δmks*<br>1,812-L |

| | | |
|---|---|---|
| IVGB248 [PAO1 parST1-PA1428(2,302-L) tetO-PA4457(1,275-L)] | (Vallet-Gely and Boccard, 2013) | Δ*mks*<br>1,275-L |
| IVGB252 [PAO1 parST1-PA3133(2,000-R) tetO-PA3573(1,509-R)] | (Vallet-Gely and Boccard, 2013) | Δ*mks*<br>1,509-R |
| IVGB253 [PAO1 parST1-PA3133(2,000-R) tetO-PA4027(1,006-R)] | (Vallet-Gely and Boccard, 2013) | Δ*mks*<br>1,006-R |
| IVGB287 [PAO1 parST1-PA0572(628-R) tetO-PA0981(1,812-L)] | (Vallet-Gely and Boccard, 2013) | Δ*mks*<br>1,812-L |
| IVGB288 [PAO1 parST1-PA4457(1,275-L) tetO-PA4027(1,006-R)] | (Vallet-Gely and Boccard, 2013) | Δ*mks*<br>1,275-L & 1,006-R |
| IVGB289 [PAO1 parST1-PA5480(92-L) tetO-PA4457(1,275-L)] | (Vallet-Gely and Boccard, 2013) | Δ*mks*<br>1,275-L |
| IVGB290 [PAO1 parST1-PA5480(92-L) tetO-PA0981(1,812-L)] | (Vallet-Gely and Boccard, 2013) | Δ*mks*<br>1,812-L |
| IVGB293 [PAO1 parST1-PA0069(82-R) tetO-PA4027(1,006-R)] | (Vallet-Gely and Boccard, 2013) | Δ*mks*<br>1,006-R |
| IVGB294 [PAO1 parST1-PA0069(82-R) tetO-PA3573(1,509-L)] | (Vallet-Gely and Boccard, 2013) | Δ*mks*<br>82-R & 1,509-L |
| IVGB296 [PAO1 parST1-PA2319(2,957-L) tetO-PA3573(1,509-R)] | (Vallet-Gely and Boccard, 2013) | Δ*mks*<br>1,509-R &2,957-L |
| IVGB297 [PAO1 parST1-PA2319(2,957-R) tetO-PA4027(1,006-R)] | (Vallet-Gely and Boccard, 2013) | Δ*mks*<br>1,006-R & 2,957-R |
| IVGB624[PAO1 *mksEF*$^{IN}$ parST1-PA2319(2,957-R)] | This paper | WT<br>2,857-R |
| IVGB625 [PAO1 *mksEF*$^{IN}$ parST1-PA2127(3,090-R)] | This paper | WT<br>3,090-L |
| IVGB626 [PAO1 *mksEF*$^{IN}$ parST1-PA0069(82-R)] | This paper | WT 82-R |
| IVGB627 [PAO1 *mksEF*$^{IN}$ parST1-PA5480(92-L)] | This paper | WT<br>92-L |
| IVGB634 [PAO1 Δsmc *mksEF*$^{IN}$ parST1-PA2319(2,957-R)] | This paper | Δ*smc*<br>2,857-R |
| IVGB635 [PAO1 Δ*smc mksEF*$^{IN}$ parST1-PA2127(3,090-R)] | This paper | Δ*smc*<br>3,090-L |

| | | |
|---|---|---|
| IVGB636 [PAO1 Δ*smc mksEF^IN^* parST1-PA0069(82-R)] | This paper | Δ*smc* 82-R |
| IVGB637 [PAO1 Δ*smc mksEF^IN^* parST1-PA5480(92-L)] | This paper | Δ*smc*<br>92-L |
| IVGB642 [PAO1 Δ*smc mksEF^IN^* parST1-PA4027(1,006-R)] | This paper | Δ*smc*<br>1,006-R |
| IVGB644 [PAO1 Δ*parS mksEF^IN^* parST1-PA2319(2,957-R)] | This paper | Δ*parS*<br>2,857-R |
| IVGB645 [PAO1 Δ*parS mksEF^IN^* parST1-PA2127(3,090-R)] | This paper | Δ*parS*<br>3,090-L |
| IVGB646 [PAO1 Δ*parS mksEF^IN^* parST1-PA0069(82-R)] | This paper | Δ*parS* 82-R |
| IVGB647 [PAO1 Δ*parS mksEF^IN^* parST1-PA5480(92-L)] | This paper | Δ*parS*<br>92-L |
| IVGB679[PAO1 Δ*parS mksEF^IN^* Δ*smc* parST1-PA0069(82-R)] | This paper | Δ*parS* Δ*smc*<br>82-R |
| IVGB701 [PAO1 Δ*smc mksEF^IN^* parST1-PA0981(1,812-L)tetO-PA3573(1,509-R)] | This paper | Δ*smc*<br>1,812-L & 1,509-R |
| IVGB702 [PAO1 Δ*smc mksEF^IN^* parST1-PA0981(1,812-L)] tetO-PA3133(2,000-R) | This paper | Δ*smc*<br>1,812-L |
| IVGB703 [PAO1 Δ*smc mksEF^IN^* parST1-PA4457(1,275-L) tetO-PA3573(1,509-R)] | This paper | Δ*smc*<br>1,275-L & 1,509-R |
| IVGB704 [PAO1 Δ*smc mksEF^IN^* parST1-PA4457(1,275-R) tetO-PA4027(1,006-R)] | This paper | Δ*smc*<br>1,275-L & 1,006-R |
| IVGB705 [PAO1 *mksEF^IN^* parST1-PA0981(1,812-L) tetO-PA3573(1,509-R)] | This paper | WT<br>1,509-R & 1,812-L |
| IVGB706 [PAO1 *mksEF^IN^* parST1-PA0981(1,812-L) tetO-PA3133(2,000-R)] | This paper | WT<br>1,812-L |
| IVGB707 [PAO1 *mksEF^IN^* parST1-PA4457(1,275-R) tetO-PA3573(1,509-R)] | This paper | WT<br>1,275-L & 1,509-R |
| IVGB708 [PAO1 *mksEF^IN^* parST1-PA4457(1,275-R) tetO-PA4027(1,006-R)] | This paper | WT<br>1,006-R & 1275-L |
| IVGB709 [PAO1 Δ*parS mksEF^IN^* parST1-PA0981(1,812-L) tetO-PA3573(1,509-R)] | This paper | Δ*parS*<br>1,509-R & 1,812-L |

| | | |
|---|---|---|
| IVGB710 [PAO1 Δ*parS mksEF^IN^* parST1-PA0981(1,812-L) tetO-PA3133(2,000-R)] | This paper | Δ*parS* 1,812-L & 2,000-R |
| IVGB711 [PAO1 Δ*parS mksEF^IN^* parST1-PA4457(1,275-R) tetO-PA3573(1,509-R)] | This paper | Δ*parS* 1,275-L & 1,509-R |
| IVGB712 [PAO1 Δ*parS mksEF^IN^* parST1-PA4457(1,275-R) tetO-PA4027(1,006-R)] | This paper | Δ*parS* 1,275-L & 1,006-R |
| IVGB754 [PAO1 *mksEF^IN^* Δ*parA* parST1-PA2319(2,957-R)] | This paper | Δ*parA* 2,857-R |
| IVGB755 [PAO1 *mksEF^IN^* Δ*parA* parST1-PA2127(3,090-R)] | This paper | Δ*parA* 3,090-L |
| IVGB756 [PAO1 *mksEF^IN^* Δ*parA* parST1-PA0069(82-R)] | This paper | Δ*parA* 82-R |
| IVGB757 [PAO1 *mksEF^IN^* Δ*parA* parST1-PA5480(92-L)] | This paper | Δ*parS* 92-L |
| IVGB840 [PAO1 Δ*parS mksEF^IN^* parST1-PA5480(92-L) tetO-PA4457(1,275-R)] | This paper | Δ*parS* 1,275-L & 92-L |
| IVGB843 [PAO1 *mksEF^IN^* parST1-PA5480(92-L) tetO-PA4457(1,275-R)] | This paper | WT 1,275-L & 92-L |
| IVGB855 [PAO1 ΔparS *mksEF^IN^* parST1-PA5480(92-L) tetO-PA4027(1,006-R)] | This paper | Δ*parS* 1,006-R & 92-L |
| IVGB856 [PAO1 *mksEF^IN^* parST1-PA5480(92-L) tetO-PA4027(1,006-R)] | This paper | WT 1,006-R & 92-L |
| IVGB923 [PAO1 *oriC ins1* parST1-PA5480(92-L) tetO-PA4027(1,006-R)] | This paper | *oriC ins1* Δ*mks* 92-L & 1006-R |
| IVGB931 [PAO1 *oriC ins1 mksEF^IN^* parST1-PA5480(92-L) post pFLP2 tetO-PA4027(1,006-R)] | This paper | *oriC ins1* 92-L & 1006-R |
| VLB114 [PAO1 Δ*smc* parST1-PA2319(2,957-R)] | This paper | Δ*mks* Δ*smc* 2,857-R |
| VLB138 [PAO1 Δ*smc* parST1-PA4027(1,006-R) tetO-PA4457(1,275-L)] | This paper | Δ*mks* Δ*smc* 1,006-R & 1,275-L |
| VLB156 [PAO1 Δ*smc* parsT1- PA3573(1,509-R) tetO-PA0981(1,812-L)] | This paper | Δ*mks* Δ*smc* 1,509-R & 1,812-L |

| | | |
|---|---|---|
| VLB157 [PAO1 Δ*smc* parST1-PA0981(1,812-L) tetO-PA4027(1,006-R)] | This paper | Δ*mks* Δ*smc* 1,006-R & 1,812-L |
| VLB158 [PAO1 Δ*smc* parST1-PA3573(1,509-R) tetO-PA4457(1,275-L)] | This paper | Δ*mks* Δ*smc* 1,509-R & 1,275-L |
| VLB21 [PAO1 Δ*parS* parST1-PA2319(2,957-L) tetO-PA0069(82-R)] | This paper | Δ*parS* Δ*mks* 82-R & 2,957-L |
| VLB40 [PAO1 Δ*smc* parST1-PA2127(3,090-L) tetO-PA0069(82-R)] | This paper | Δ*mks* Δ*smc* 82-R 3,090-L |
| VLB47 [PAO1 Δ*smc* parST1-PA2127(3,090-R) tetO-PA5480(92-L)] | This paper | Δ*mks* Δ*smc* 92-L & 3,090-L |
| Oligonucleotides | | |
| Adapters | Marbouty et al 2015 | N/A |
| Software and Algorithms | | |
| Bowtie2 | (Langmead and Salzberg, 2012) | http://bowtie-bio.sourceforge.net/bowtie2/index.shtml |
| Matlab | The MathWorks Inc | https://fr.mathworks.com/products/matlab.html |
| Pipeline to analyze 3C-seq data | (Lioy et al., 2018) | https://github.com/koszullab/E_coli_analysis |
| Plasmids | | |
| pPSV35Ap-TetR-Cfp-yGfp-ParBT1 | (Vallet-Gely and Boccard, 2013) | N/A |
