## Supplementary Figures for "Distinct activities of bacterial condensins for chromosome management in *Pseudomonas aeruginosa*"

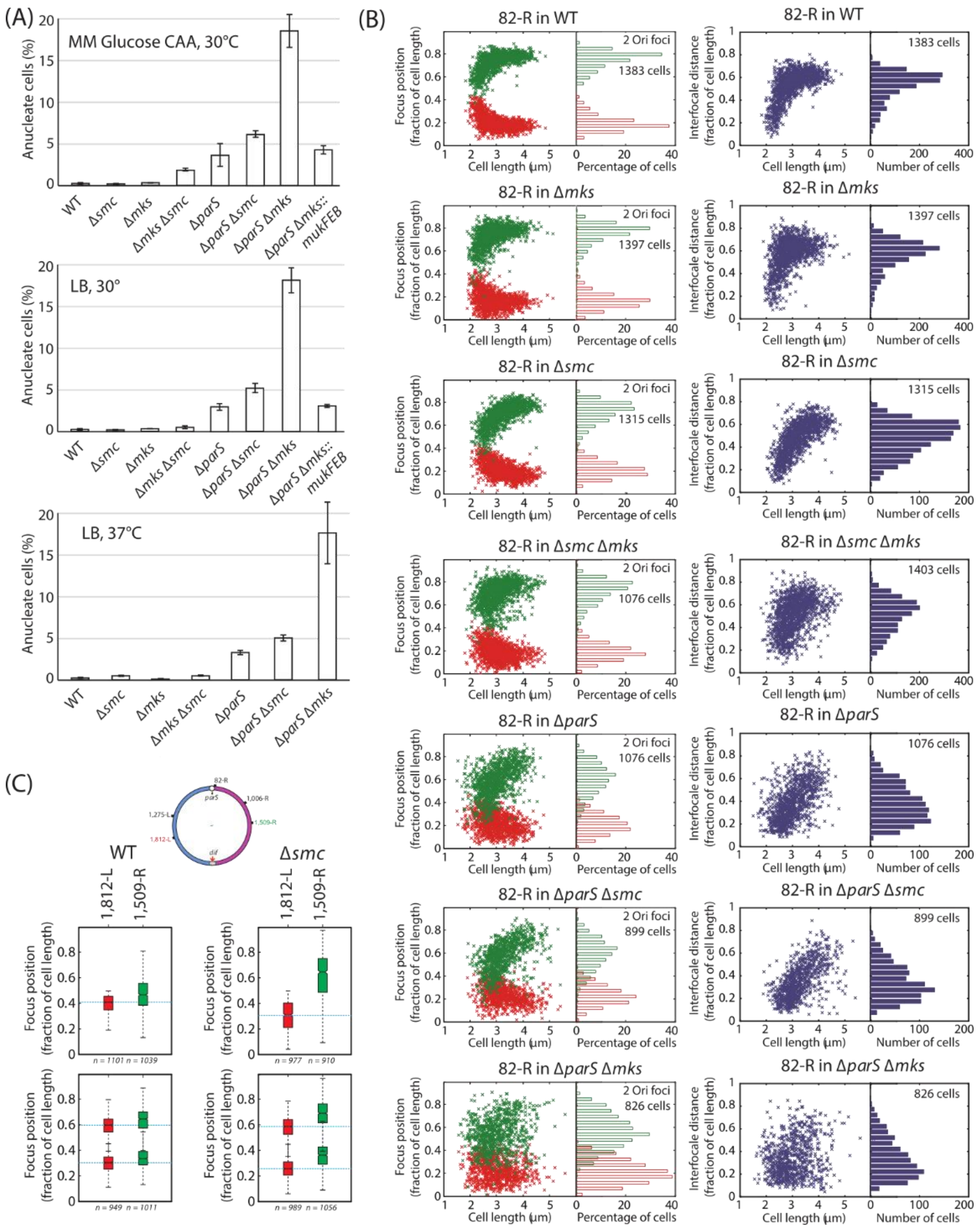


**Supplementary Figure 1. Differential impact of ParABS, MksBEF and Smc-ScpAB on chromosome segregation and positioning. (A)** Percentage of anucleate cells (white bars) in cultures of the different mutants grown in minimal medium supplemented with glucose and casamino acids at 30°C (left panel), in LB at 30°C (middle panel) and in LB at 37°C (right panel). Histograms and error bars represent the mean and standard deviation for at least three independent experiments. **(B)** (Left) Position of the two copies of a chromosomal locus located near the *parS* sites (82-R) in two-foci cells, in different genetic backgrounds (indicated above each graph). The relative position of the two copies of the fluorescent tag inside the cell are represented according to cell length. Cells are arbitrarily oriented, with the 0 pole being the one closest to a fluorescent tag. (Right) Distance between the two copies of a chromosomal locus located near the *parS* sites (82-R) in two-foci cells (whose position are represented in Figure 1D), in different genetic backgrounds (indicated above each graph). The relative distance is represented according to cell length. **(C)** Schematic representation of the position of the chromosomal loci studied. The two chromosomal loci whose relative position inside the cells are represented in **(D)** are highlighted in green and red. **(D)** Top panels: relative position inside the cells of two chromosomal loci located at similar distance from *oriC* (highlighted in green and red in (C)), in the wild type strain (left panels) and in the Δ*smc* mutant (right panels). Boxplot representations are used, indicating the median (horizontal bar), the 25th and the 75$^{th}$ percentile (open box) and the rest of the population except for the outliers (whiskers). Outliers are defined as 1.5×*IQR* or more above the 75$^{th}$ percentile or 1.5×*IQR* or more below the first 25th percentile quartile. Cells are arbitrarily oriented, the 0 pole being the one closest to the 1,812-L locus. Experiments have been performed at least twice independently; one representative example is shown here. Bottom panels: interfocal distances (in micrometers) between the two chromosomal loci, in cells containing one copy of each locus (top panel) or two copies of each locus (bottom panel). The percentage of cells in which the distance is of a certain value is plotted, for the wild type strain (white) and the Δ*smc* mutant (yellow). Histograms and error bars represent the mean and standard deviation for two independent experiments.

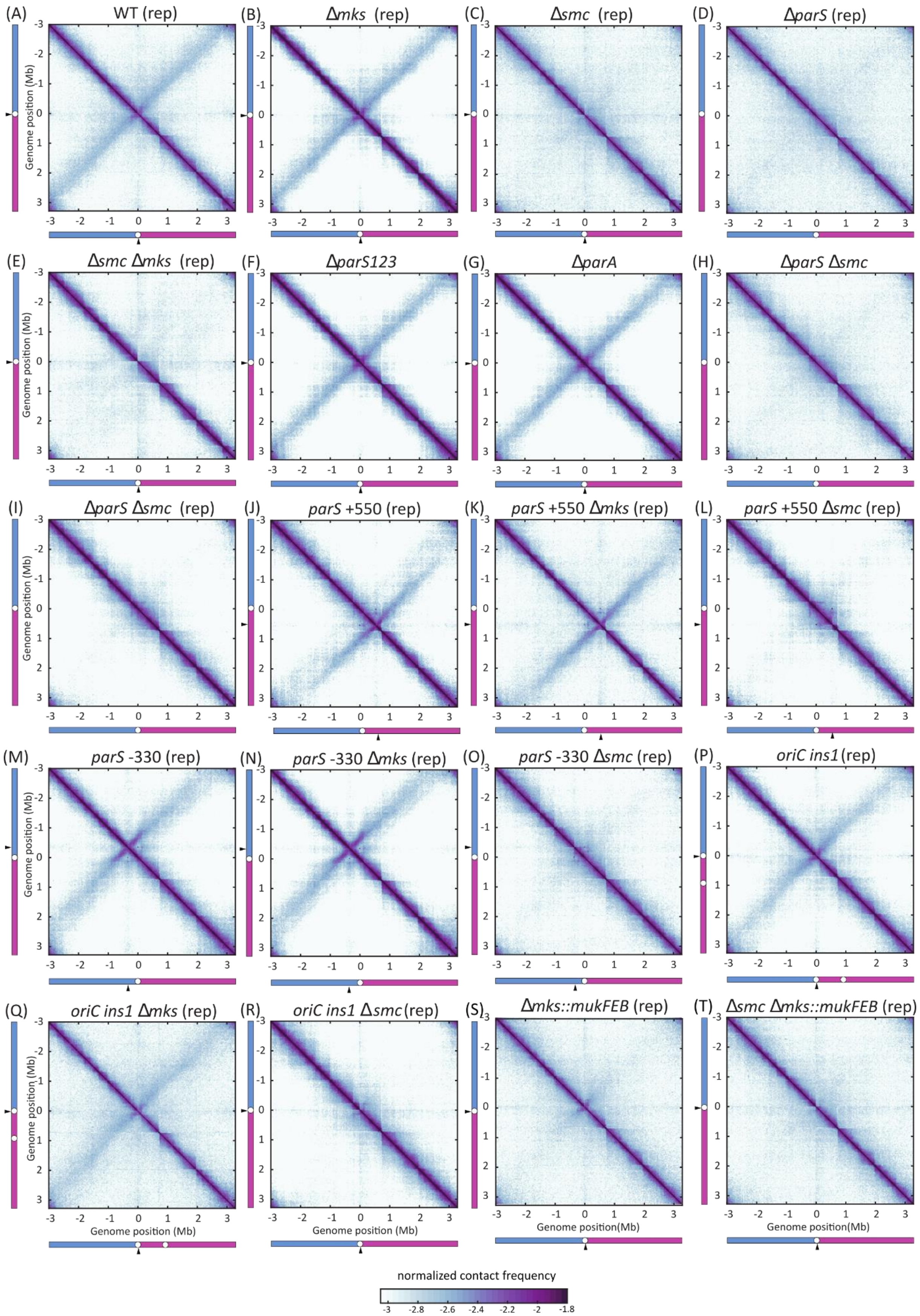

(A) WT (rep)
(B) Δmks (rep)
(C) Δsmc (rep)
(D) ΔparS (rep)
(E) Δsmc Δmks (rep)
(F) ΔparS123
(G) ΔparA
(H) ΔparS Δsmc
(I) ΔparS Δsmc (rep)
(J) parS +550 (rep)
(K) parS +550 Δmks (rep)
(L) parS +550 Δsmc (rep)
(M) parS -330 (rep)
(N) parS -330 Δmks (rep)
(O) parS -330 Δsmc (rep)
(P) oriC ins1 (rep)
(Q) oriC ins1 Δmks (rep)
(R) oriC ins1 Δsmc (rep)
(S) Δmks::mukFEB (rep)
(T) Δsmc Δmks::mukFEB (rep)
Genome position (Mb)
normalized contact frequency
-3
-2.8
-2.6
-2.4
-2.2
-2
-1.8

**Supplementary Figure 2: Normalized contact maps obtained for different strains grown in minimal medium supplemented with citrate at 30°C.** Abscissa and ordinate axis represent genomic coordinates. The colorscale reflects the frequency of contacts between two regions of the genome, from white (rare contacts) to dark purple (frequent contacts). Maps obtained for the wild type strain **(A)**, the Δ*mks* mutant **(B)**, the Δ*smc* mutant **(C)**, the Δ*parS* mutant **(D)**, the Δ*smc* Δ*mks* mutant **(E)**, the Δ*parS*123 mutant **(F)**, the Δ*par*A mutant **(G)**, the Δ*parS* Δ*smc* mutant **(H and I),** the *parS* $^{+550}$ **(J)**, *parS* $^{+550}$ Δ*mks* **(K)**, *parS* $^{+550}$ Δ*smc* **(L)**, *parS* $^{-330}$ **(M)**, *parS* $^{-330}$ Δ*mks* **(N)**, *parS* $^{-330}$ Δ*smc* **(O),** *oriC ins1* **(P)**, *oriC ins1* Δ*mks* **(Q)**, *oriC ins1* Δ*smc* **(R)** Δ*mks::mukFEB* **(S)** and Δ*smc* Δ*mks::mukFEB* **(T)** strains.

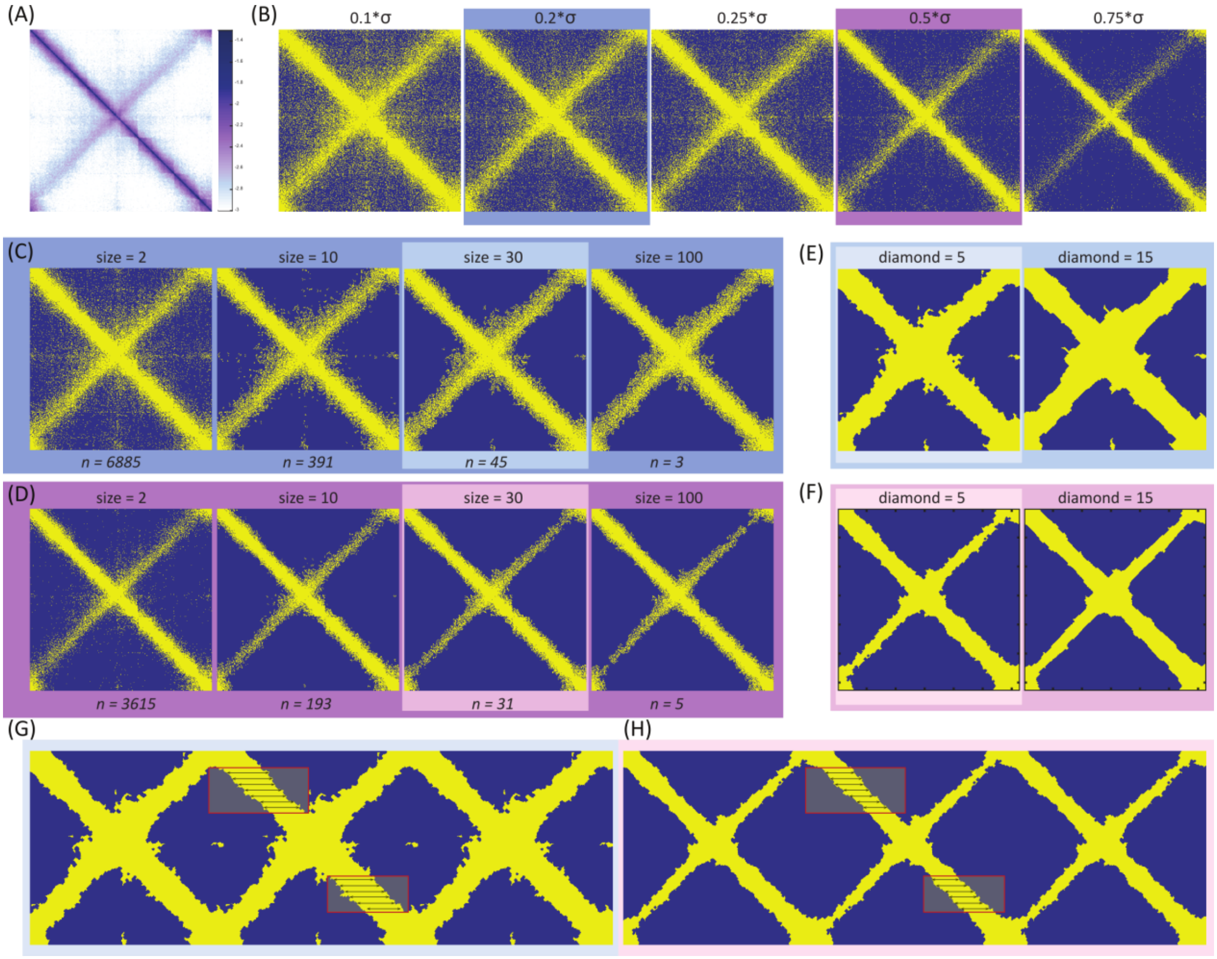


**Supplementary Figure 3: Illustration of the three step process used to determine the width of the primary diagonal, i.e. the range of *cis* contacts of chromosomal loci. (A)** Contact map to be analyzed. **(B)** Binary contact maps obtained after setting the threshold for significant interactions at 0.1, 0.2, 0.25, 0.5 and 0.75 times the standard deviation (defined as σ= 1.4826*mad (median absolute deviation)) above the median. Contact frequencies above or below the threshold were assigned a value of 1 or 0, respectively, generating a binary contact map in which significant interactions between chromosomal loci are represented in yellow whereas non-significant interactions are represented in blue. Abscissa and ordinate axis represent genomic coordinates. **(C)** and **(D)** Binary contact maps obtained by varying the size of the connected element considered as significant (smaller elements are discarded as non-significant background noise). (C) starting point is the 0.2*σ map whereas (D) starting point is the 0.75*σ one, as indicated by the colored area surrounding the maps. Yellow dots indicate a significant interaction whereas blue dots indicate a non-significant interaction. Abscissa and ordinate axis represent genomic coordinates. **(E)** and **(F)** Binary contact maps obtained using different sizes of the diamond shape used to fill out the empty points comprised in the connected elements. As indicated by the colored area surrounding the maps, (E) comes from (C) size = 30 and (F) comes from (D) size = 30. **(G)** and **(H)** Schematic representations of the measurement of the range of *cis* contacts for chromosomal loci belonging to chromosome arms. Region in which significant interactions were measured for each chromosomal locus (black arrows) are represented as red rectangles.

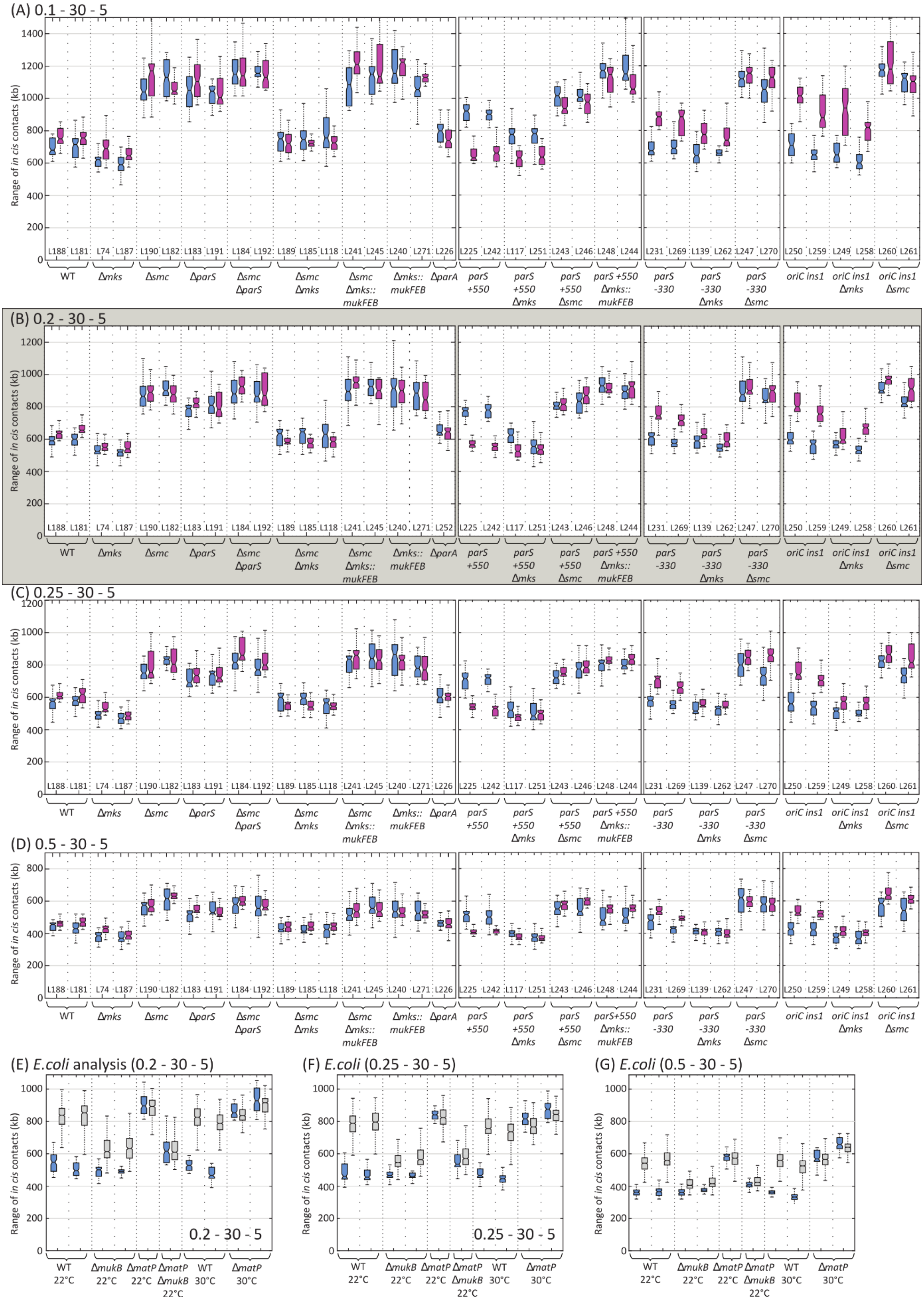

(A) 0.1 - 30 - 5
Range of in cis contacts (kb)
0 200 400 600 800 1000 1200 1400
L188 L181 L74 L187 L190 L182 L183 L191 L184 L192 L189 L185 L118 L241 L245 L240 L271 L226 L225 L242 L117 L251 L243 L246 L248 L244 L231 L269 L139 L262 L247 L270 L250 L259 L249 L258 L260 L261
WT
Δmks
Δsmc
ΔparS
Δsmc ΔparS
Δsmc Δmks
Δsmc Δmks:: mukFEB
Δmks:: mukFEB
ΔparA
parS +550
parS +550 Δmks
parS +550 Δsmc
parS +550 Δmks:: mukFEB
parS -330
parS -330 Δmks
parS -330 Δsmc
oriC ins1
oriC ins1 Δmks
oriC ins1 Δsmc
(B) 0.2 - 30 - 5
Range of in cis contacts (kb)
0 200 400 600 800 1000 1200
L188 L181 L74 L187 L190 L182 L183 L191 L184 L192 L189 L185 L118 L241 L245 L240 L271 L252 L225 L242 L117 L251 L243 L246 L248 L244 L231 L269 L139 L262 L247 L270 L250 L259 L249 L258 L260 L261
(C) 0.25 - 30 - 5
Range of in cis contacts (kb)
0 200 400 600 800 1000 1200
L188 L181 L74 L187 L190 L182 L183 L191 L184 L192 L189 L185 L118 L241 L245 L240 L271 L226 L225 L242 L117 L251 L243 L246 L248 L244 L231 L269 L139 L262 L247 L270 L250 L259 L249 L258 L260 L261
(D) 0.5 - 30 - 5
Range of in cis contacts (kb)
0 200 400 600 800
L188 L181 L74 L187 L190 L182 L183 L191 L184 L192 L189 L185 L118 L241 L245 L240 L271 L226 L225 L242 L117 L251 L243 L246 L248 L244 L231 L269 L139 L262 L247 L270 L250 L259 L249 L258 L260 L261
(E) E.coli analysis (0.2 - 30 - 5)
(F) E.coli (0.25 - 30 - 5)
(G) E.coli (0.5 - 30 - 5)
Range of in cis contacts (kb)
0 200 400 600 800 1000
0.2 - 30 - 5
0.25 - 30 - 5
WT 22°C
ΔmukB 22°C
ΔmatP 22°C
ΔmatP ΔmukB 22°C
WT 30°C
ΔmatP 30°C

**Supplementary Figure 4: (A-D) Quantification of the range of *cis* contacts for all contact maps of this study and for *E. coli* contact maps of the wild type strain, the Δ*matP* and the Δ*mukB* mutants.** Using different thresholds for a significant interaction: 0.1*σ **(A)**, 0.2*σ **(B)**, 0.25*σ **(C)** and 0.5*σ **(D)**. Boxplot representations are used, indicating the median (horizontal bar), the 25th and the 75$^{th}$ percentile (open box) and the rest of the population except for the outliers (whiskers). Outliers are defined as 1.5×*IQR* or more above the 75$^{th}$ percentile or 1.5×*IQR* or more below the first 25th percentile quartile. The 0.2*σ threshold is highlighted in grey as it is the threshold chosen for the main figures. **(E-G)** Quantification of the range of *cis* contacts for *E. coli* contact maps of the wild type strain, the Δ*matP* and the Δ*mukB* mutants, previously published in Lioy et al., 2018. Strains were grown in Minimal medium supplemented with glucose and casamino acids, at the temperature indicated below the graphs with the genetic background. Different thresholds were used, 0.2*σ **(E)**, 0.25*σ **(F)** and 0.5*σ **(G)**.

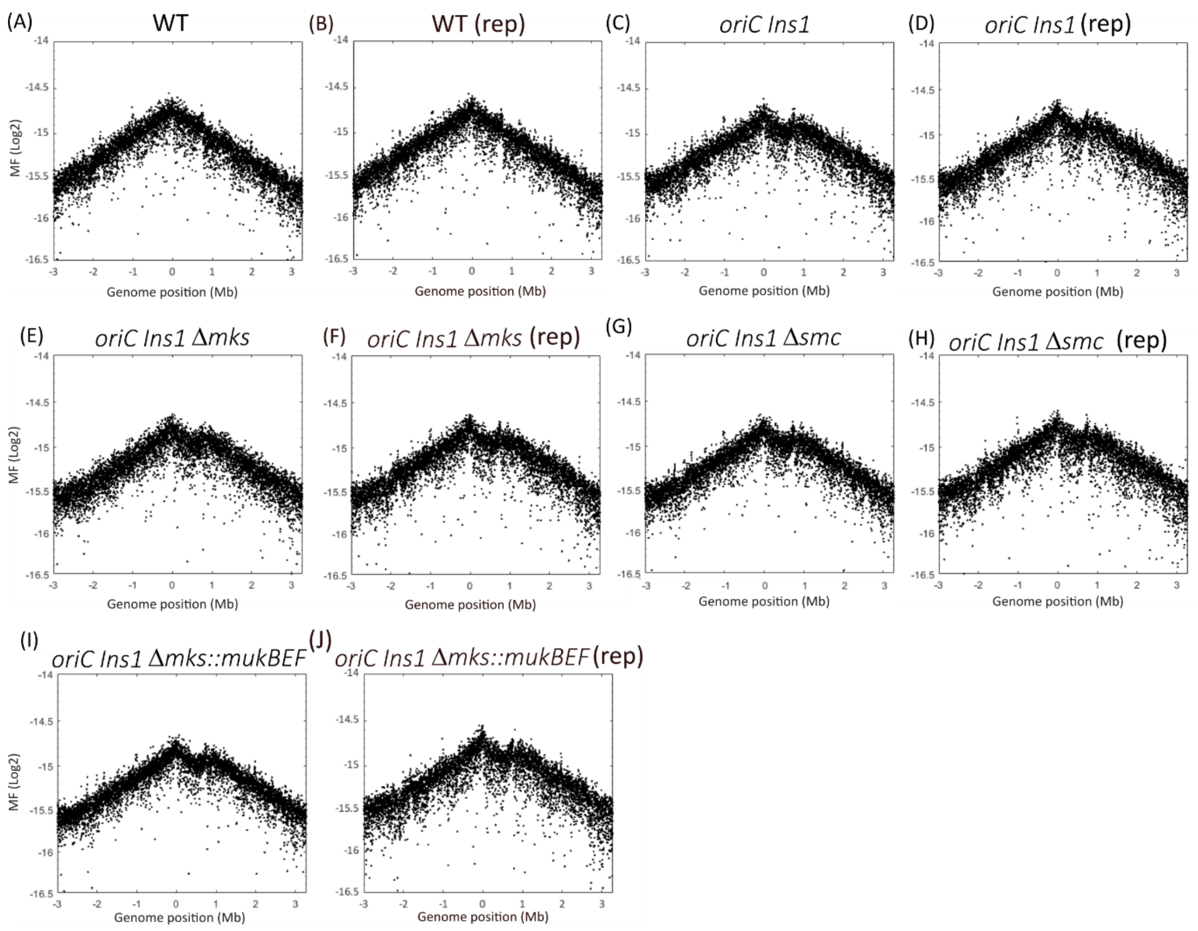


**Supplementary Figure 5: Marker frequency analyses for different strains grown in minimal medium supplemented with citrate at 30°C.** Marker frequencies for WT (A) and (B), *oriC ins1* **(C) and (D)**, *oriC ins1* Δ*mks* **(E)**, *oriC ins1* Δ*smc* **(R)** Δ*mks::mukFEB* **(S)** and Δ*smc* Δ*mks::mukFEB* **(T)** strain marker frequencies (black dots after trimming and normalisation on the total number of reads, see STAR methods) are represented in Log2 as a function of the genome position.

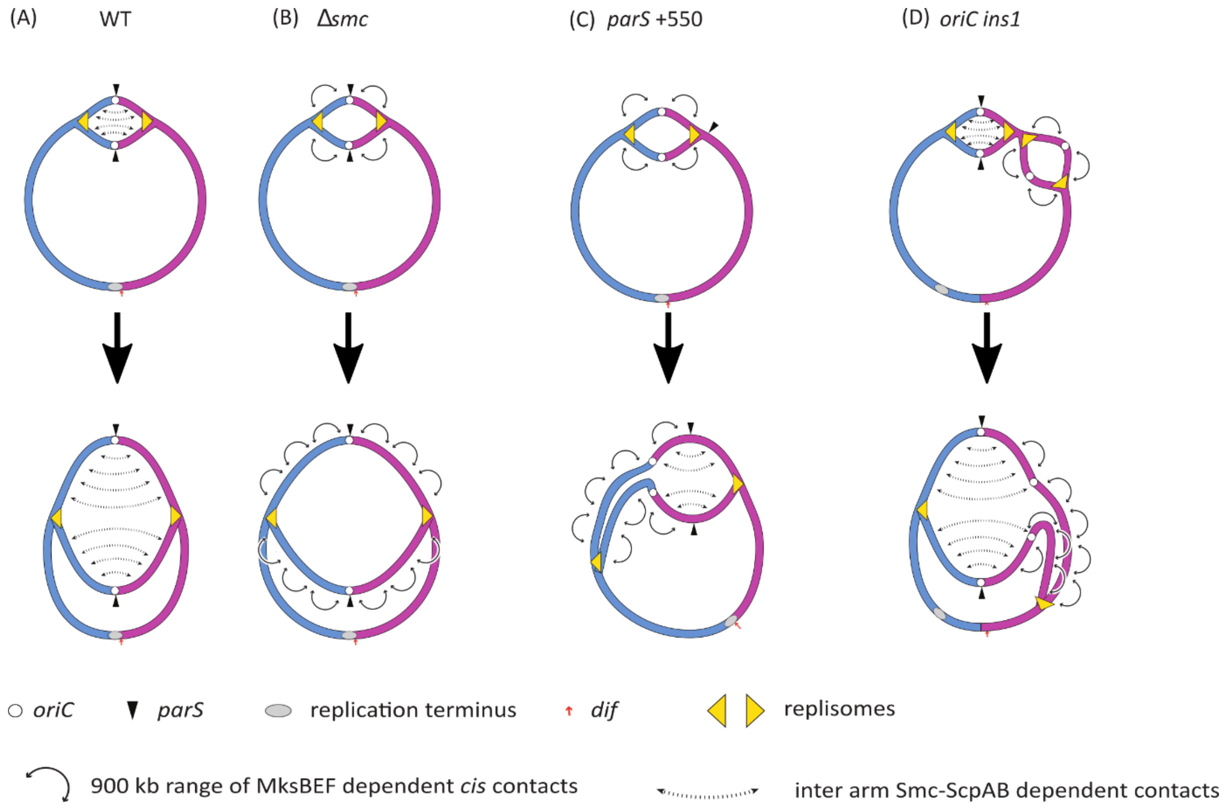


**Supplementary Figure 6: Recapitulative schematization of the different chromosomal contacts observed in *P. aeruginosa* according to different chromosome configurations during the replication cycle**. Overall contacts in the WT strain **(A)**, in the Δsmc mutant **(B)**, in the strain with the displaced *parS* site (*parS* $^{+550}$) **(C)** and in the strain with the additional ectopic *oriC* (*oriC ins1*) **(D).**
